# Systemic Lupus Erythematosus Serum Stimulation of Human Intestinal Organoids Induces Changes in Goblet Cell Differentiation and Mitochondrial Fitness

**DOI:** 10.1101/2023.07.04.547690

**Authors:** Inga Viktoria Hensel, Szabolcs Éliás, Michelle Steinhauer, Bilgenaz Stoll, Salvatore Benfatto, Wolfgang Merkt, Stefan Krienke, Hanns-Martin Lorenz, Jürgen Haas, Brigitte Wildeman, Martin Resnik-Docampo

## Abstract

Human intestinal epithelial cells are the interface between potentially harmful luminal content and basally residing immune cells. Their role is not only nutrient absorption but also the formation of a tight monolayer that constantly secrets mucus creating a multi-layered protective barrier. Alterations in this barrier can lead to increased gut permeability which is frequently seen in individuals with chronic extraintestinal autoimmune diseases, such as Systemic Lupus Erythematosus (SLE). Despite recent advances in identifying alterations in gut microbiota composition in SLE patients, not much attention has been given to the epithelial barrier itself. To date, it remains largely unexplored which role and function intestinal epithelial cells have in SLE pathology. Here, we present a unique near-physiologic *in vitro* model specifically designed to examine the effects of SLE on the epithelial cells. We utilize human colon organoids that are stimulated with serum obtained from SLE patients. Combining bulk and scRNA transcriptomic analysis with functional assays revealed that SLE serum stimulation induced a unique expression profile marked by a type I interferon gene signature. Additionally, organoids exhibited decreased mitochondrial fitness, alterations in mucus composition and imbalanced cellular composition. Similarly, transcriptomic analysis of SLE human colon biopsies revealed a downregulation of epithelial secretory markers. Our work uncovers a crucial connection between SLE and intestinal homeostasis that might be promoted *in vivo* through the blood, offering insights into the causal connection of barrier dysfunction and autoimmune diseases.

## Introduction

The intestine has a substantial surface area that harbors the highest quantity of immune cells in close proximity to the tremendous number of microbes in our body.^1–3^ Thus, a functional epithelial barrier is crucial for separating these two compartments to ensure intestinal homeostasis and overall health. Under physiological conditions this function is maintained by constant renewal of the barrier containing a balanced cellular composition.^4–6^ Intestinal stem cells and transit-amplifying (TA) cells proliferate and give rise to different specialized cells committed to either absorptive or secretory lineage, represented by the two major cell types: colonocytes and goblet cells. While colonocytes mainly contribute to maintaining fluid balance and absorbing nutrients, goblet cells are considered as key players of mucosal barrier integrity. The secreted mucus forms a layer which is not only essential as defense against microbial infiltration, but also acts as a niche for commensals.^5, 7–9^ Its composition is highly dynamic, can be influenced not only by the abundance of mucus core proteins and antimicrobial peptides secreted to it, but also by factors such as ion concentration, pH, and hydration state^10, 11^. The cell type composition and therewith barrier function is continuously influenced by signals from neighboring immune and stromal cells, cytokines, bacterial metabolites, and nutrients. Hallmarks of pathological conditions like inflammatory bowel disease and infections are shifts in cell type proportions and alterations in mucus composition affecting barrier function and increasing gut permeability.^12, 13^ Recent research advancements have started to shed light on the broader implications of gut permeability. The discovery that the intestine plays a considerable role in the pathogenesis of a number of extraintestinal systemic diseases marks a paradigm shift, expanding our understanding of the pivotal role the intestine plays in overall health and disease^14, 15^.

Systemic lupus erythematosus (SLE), a multifaceted autoimmune disease which is characterized by systemic inflammation affecting skin, kidney and the central nervous system.^16^ Recent reports show that SLE patients have an altered microbiome^17–22^ and increased intestinal permeability^15, 23–26^. Limited knowledge exists regarding the role and contribution of the intestine in the pathogenesis and progression of the disease. Especially in which extent the intestinal epithelial cells are affected remains to be elucidated. SLE manifests with a dysregulation of the immune system and is characterized by elevated type I interferon levels and presence of autoantibodies in serum.^27–30^ The blood is a complex body fluid that serves as transport medium to supply all cells of the body with nutrients and oxygen. The intestine is the most intensely perfused organ. Especially the metabolically highly active epithelial cells are in close contact with the underlying vasculature ensuring swift exchange and efficient nutrient absorption necessary for homeostasis.^31^ However, this may also facilitate an influx of circulating cytokines and autoantibodies contained in the blood to the intestine with the potential to alter epithelial cell dynamics and barrier function^32–34^. A comprehensive understanding of this interconnection is of great importance to elucidate the role of the intestinal epithelial barrier in autoimmune diseases and for the development of novel therapeutic approaches.

In order to fully understand this complex interplay in a multicellular environment, it is necessary to develop a model that can explore the impact of blood components on the intestinal epithelium in near physiological conditions. This model, however, should be strategically designed to only include intestinal epithelial cells. By specifically focusing on these cells, we can eliminate potential interference from other components on opposite sides of the barrier. This includes microbiota on the luminal side, as well as immune and stromal cells on the basolateral side. In this way, we can isolate and observe the direct effects of blood components on intestinal epithelial cells, providing us with a more controlled and precise understanding of these interactions. To this end, the development of organoids as advanced primary cell models has revolutionized the design of human tissue models. These self-organized, three-dimensional structures, derived from adult stem cells, surpass traditional cell models while recapitulating the organ architecture and plasticity found *in vivo*.^35^Organoids have become a notable alternative to animal models, offering a more reliable and human-relevant approach for addressing long-standing medical challenges. The relevance of organoids as a model system has been recognized by the FDA that recently approved non-animal models in the drug development process.^36^ This marks the beginning of a new era in biomedical research. Our research takes a pioneering approach by integrating serum derived from SLE patients into intestinal organoids enabling us to simulate the impact of blood components on the intestinal epithelium while minimizing the interference caused by the intricate intestinal cell milieu, providing a cleaner, more controlled environment for our explorations and analyses.

Herein, we report that our organoid model contains all relevant cell types found in colon. We show that SLE serum stimulation can induce alterations in the expression profile, which is dependent on type I interferon signaling, induced through a synergistic effect of all serum contained factors and is specific to SLE. We can show that the secretory lineage is majorly affected, and the mucus layer potentially weakened by alterations in mucus composition, specifically of antimicrobial peptides. A similar impact on the secretory lineage and mucus layer is also observed in Ulcerative Colitis suggesting epithelial barrier dysfunction as hallmark in SLE pathogenesis. Conclusively, our *in vitro* model emulates a pathological condition where a complex mix of cytokines and other SLE specific factors secreted at the site of inflammation and distributed systemically reaching the highly perfused intestine via the blood can impact the epithelial barrier. This innovative disease model holds the potential to unveil unique insights into disease mechanisms that would otherwise be challenging to investigate. Additionally, it holds the potential as tool to identify innovative treatment approaches and points of intervention.

## Results

### 1. Organoids show SLE specific response to serum stimulation

To understand the effect of SLE on the intestinal epithelial barrier, we generated organoid line I and II from the descending colon of two healthy donors, cultured them in conditions that conserved the native cellular diversity^35^ and stimulated them for 72h with 5% serum from treatment-naїve SLE patients or sex- and age-matched controls (Fig. 1A and Table 1). Serum stimulation did not induce any significant phenotypical changes in size or shape (Fig. 1B). There was no evidence for increased cytotoxicity or apoptosis (Fig. 1C and Suppl. Fig. 1A). Expression profiles of the stimulated organoids were marked by a distinct spread in expression profiles for both stimulation conditions across two tested organoid lines (Fig. 1D). The first principal component (PC1), accounting for 39% of the total variance, displayed a separation of a subgroup of organoids stimulated with SLE sera from the controls in organoid line I. Interestingly, the same SLE sera induced an even more pronounced separation in organoid line II along the PC1 axis. This consistent separation across both organoid lines strongly suggested a donor-independent effect of SLE serum stimulation on epithelial cells. The second principal component (PC2), accounting for 15% or 21% of the total variance, further differentiated the samples within each condition, likely reflecting serum sample-specific responses.

**Figure 1:**
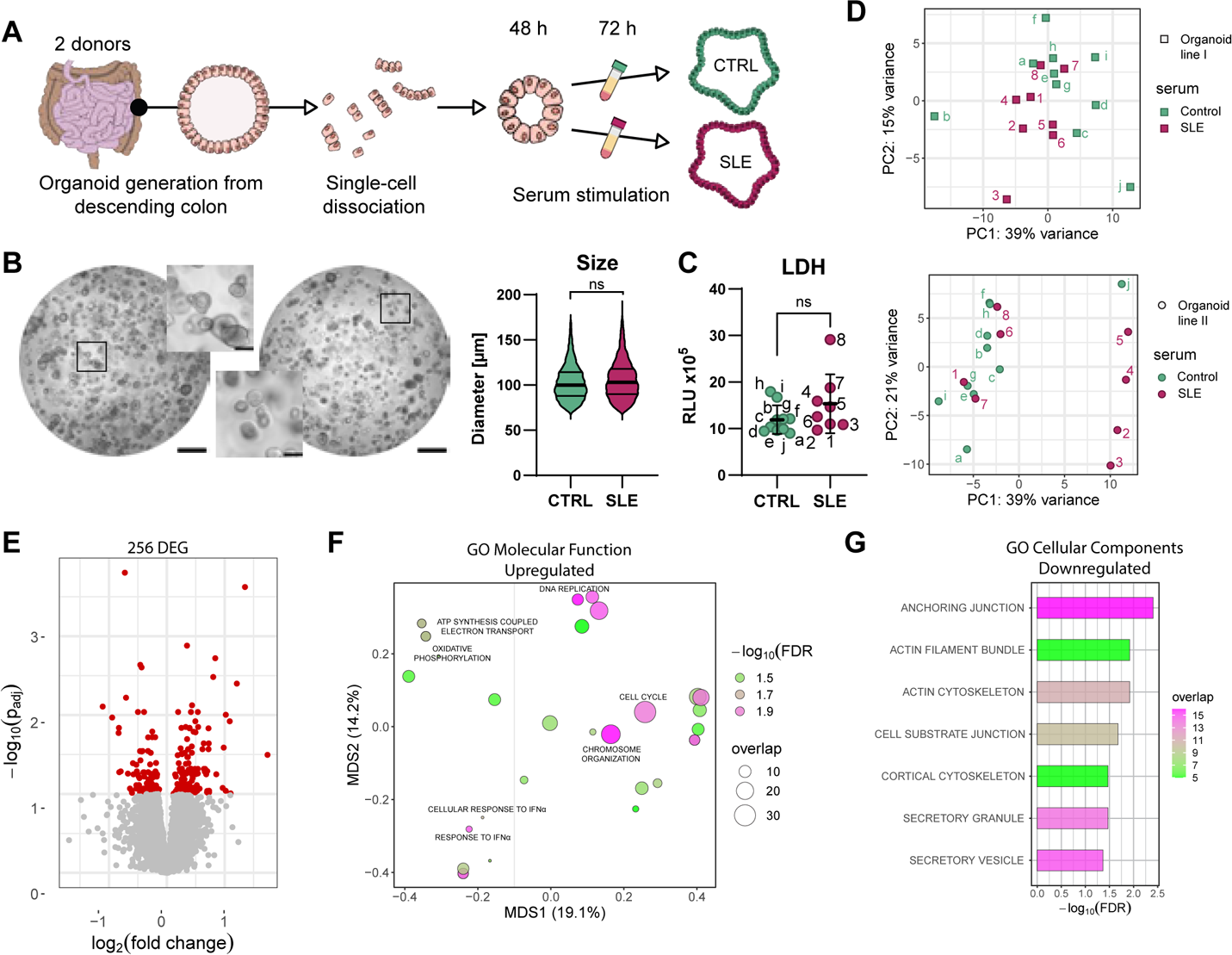
Transcriptomic profiling of serum stimulated organoids reveals SLE specific responses. (A) Schematic representation of the experimental approach. (B) Brightfield microscopy images of colonic organoids after 72h stimulation with either control serum (left) or SLE serum (right). Insets showcase enlarged areas, scale bars 1000 μm and 100 μm respectively. Graphical summary demonstrates no significant difference in organoid size between control and SLE stimulated samples; in total 10000 organoids per condition were analyzed, lines represent the quartiles and the mean (analyzed using chi-square test considering 100 averaged values). (C) Analysis of lactate dehydrogenase activity (LDH) reveals absence of apoptosis or cytotoxicity in SLE serum-stimulated organoid. Each dot corresponds to one well of organoids. Statistically significant differences were not observed, as determined by Kolmogorov-Smirnov test. (D) PCA Highlights unique responses in organoids stimulated with SLE Serum after 72h. PCA plot 1 (organoid line I) shows gene expression differences between organoids treated with control (green) and SLE (pink) serum. PC1, explaining 39% of the total variance, separates a subgroup of organoids stimulated with SLE serum. PCA plot 2 (organoid line II) reveals a more pronounced distinction in the SLE serum-stimulated organoids along the PC1 axis, reinforcing the consistent impact of SLE serum across different organoid lines. PC2, accounting for 15% and 21% of the variance in each plot respectively, provides further differentiation within each condition, likely due to the unique responses to individual serum samples. (E) Volcano Plot of bulk RNA-Seq data highlights 256 differentially expressed genes of colon organoids 72h post serum stimulation, red dots highlighting the significantly misregulated genes. (F) Gene Ontology (GO) analysis of upregulated genes reveals overrepresentation in key biological processes. (G) GO analysis indicates enrichment of specific processes among downregulated genes. Unless otherwise specified, all experiments depicted in this figure were analyzed with n= 10 control serum (samples a to j) and n=8 SLE serum (samples 1 to 8).

Taken together, these results reinforced the inherent complexity of serum stimulated organoids and the need to consider both stimulation and organoid donor-specific effects for downstream analysis and data interpretation. Therefore, we pooled the data from both organoid donors accounting for sex and individual characteristics of the organoid line. In addition, we integrated the response of different cell types given the fact that the cell type composition in both organoid lines varied due to individual proliferation and differentiation dynamics (Suppl Fig. 1B). The combined analysis resulted in 256 differentially expressed genes (Fig. 1E). Gene Ontology (GO) analysis of the upregulated genes showed an overrepresentation of terms connected to cell cycle, chromosome organization and replication as well as mitochondrial function and interferon signaling (Fig. 1F). The downregulated genes showed an enrichment of terms connected to secretion, cytoskeleton, and anchoring junctions of the cells (Fig. 1G).

Intriguingly, the overall results showed that similar pathways were altered in epithelial cells as previously only described in immune cells of SLE patients.^37^ Thus, the distinct response of organoids to SLE patient derived serum, consistent in both organoid lines, holds the potential to unravel how SLE specific mechanisms can influence the intestinal epithelial barrier.

### 2. Type I Interferon drives the expression changes induced by SLE serum

We wanted to explore the overrepresentation of terms connected to interferon (IFN) signaling more in depth. Out of the 256 DEG we found 22 genes to be connected to type-I IFN signaling (Fig. 2A). A significant majority of them were upregulated. They accounted for 32% of the 25 highest upregulated genes underlining the role of type-I IFN in the response induced by SLE serum stimulation. To investigate whether the expression of IFN related genes was altered in both organoid lines by the different patient sera, we selected a range of IFN inducible genes that were represented in several GO terms (Fig. 2B). The representation of their expression in a heatmap revealed an upregulation in all the SLE serum stimulated organoids except for two SLE sera (Fig. 2B; Suppl. Fig. 2A; Suppl. table 1). On the contrary only two of the control sera showed a similar expression as the SLE sera stimulated organoids indicating the SLE specificity of this response.

**Figure 2:**
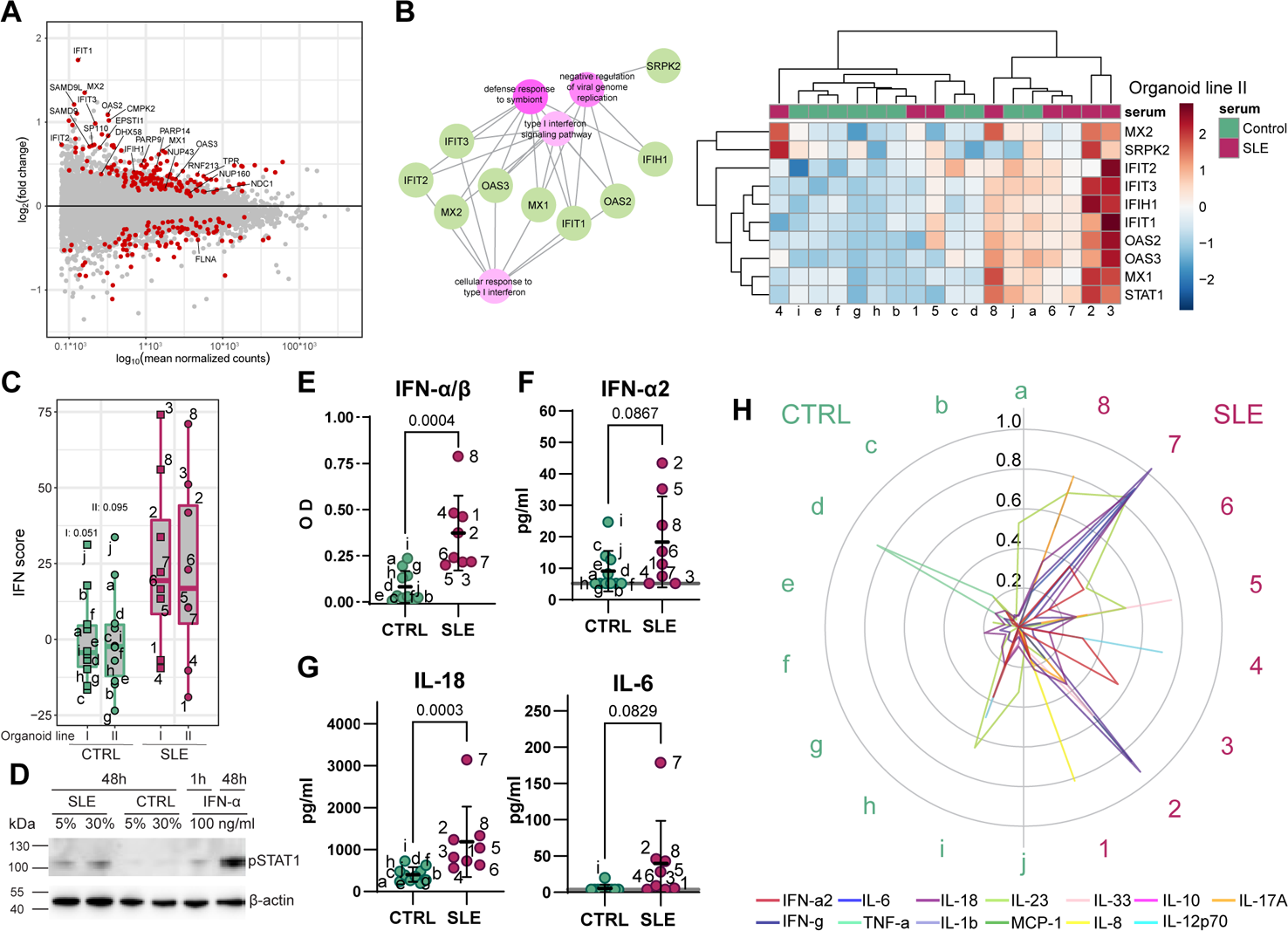
Organoid expression profile induced by complex cytokine composition in SLE serum is IFN-α driven. A: MA plot featuring 22 out of 256 differentially expressed genes associated with IFN signaling, constituting 32% of the top 25 most upregulated genes. B: Network diagram illustrating selected IFN-inducible genes interconnected with different IFN-related GO terms. Heatmap visualizing the expression of these selected IFN-inducible genes, showing an upregulation across all SLE serum-stimulated organoids from line II, except for two specific SLE sera. C: IFN score representation indicating the upregulation of IFN inducible genes in organoids stimulated by SLE serum, derived from z-scores of normalized counts, statistical differences determined by one way ANOVA with Holm-Šidák’s multiple comparisons test. D: Phosphorylation of STAT1, displaying a concentration-dependent response to SLE serum and minimal reaction in control serum-stimulated organoids after 48h. IFN-α represents the positive control after 1h and 48h of stimulation. Each sample represents two pooled wells of organoids stimulated with serum from one donor. E: Elevated type I interferon levels in SLE serum detected using an IFN-α/β reporter cell line. Each dot represents one well of serum stimulated cells (n=10 and n=8 for control or SLE serum respectively). F: IFN-α2 levels in SLE serum confirmed via multiplex ELISA, validating the increased interferon levels. Grey line shows ½ the detection limit. n=10 and n=8 for control or SLE serum respectively. G: Multiplex ELISA data highlighting elevated IL-18 and IL-6 levels in SLE patient sera compared to controls. In only one control IL-6 was detectable compared to six out of eight SLE serum samples. Half the detection limit concentration was used for calculation if the cytokine was undetectable. Grey line shows ½ the detection limit. n=10 and n=8 for control or SLE serum respectively. H: Radar plot illustrating the distinct cytokine composition in each serum sample, quantified by Multiplex ELISA. Each line color represents one cytokine, levels are shown as fraction, and normalized to the highest and lowest value for each cytokine. Green and pink labels represent control and SLE samples respectively. Unless otherwise indicated, an unpaired t-test was used for statistical analysis. Unless otherwise specified, all experiments depicted in this figure were analyzed with n= 10 control serum (samples a to j) and n=8 SLE serum (samples 1 to 8).

A common way to analyze the type-I IFN levels and activity in SLE serum samples is to analyze the expression profile of interferon signature genes (IFNSG) in SLE whole blood samples.^38^ Given the fact that we were studying epithelial cells we decided to choose the IFNSG based on IFN-α stimulated organoids. Considering only genes relevant in IFNα/β signaling we identified 27 genes which were specific for epithelial cell response to a low dose of IFN-α (Suppl. Fig. 2B). We used this panel of genes (irrespective of their significance of expression) to calculate an IFN score from the normalized counts of SLE and control serum stimulated organoids (Fig. 2C). We could see an overall higher score in organoids stimulated with SLE serum compared to the control irrespective of the organoid line. The IFN score gave us a powerful tool to assess the expression of IFNSG unique to each serum independent of the significance in expression identified by the pooled data. Thus, we were able to integrate and interpret the effect by IFN in a more robust way than just focusing on single gene expression or DEGs which are based on the mean expression. In addition, we ruled out that the seen IFN signature was caused by endogenous expressed IFN by confirming the absence of IFNα1 and IFNβ1 transcripts (Suppl. Fig. 2C). Additionally, we could confirm the activation of IFN signaling by upregulation and phosphorylation of STAT1 which was SLE serum concentration dependent and almost absent in control serum stimulated organoids (Fig. 2D, Suppl. Fig. 2D). Only with SLE serum or IFNα stimulation we could detect IFIT3, Interferon Induced Protein with Tetratricopeptide Repeats 3, abundance (Suppl. Fig. 2D).

As a next step we wanted to quantify type I IFN levels of the used serum samples. Since it is challenging to measure it in serum^39, 40^, we employed two different approaches analyzing not only the concentration but validating elevated levels by a functional read out. With an IFN-α/β reporter cell line that was stimulated with serum we could detect a significant increase of type I IFN levels in SLE serum (Fig. 2E). We then used a bead-based immunoassay to quantify IFN-α2 levels which confirmed the increased interferon levels in SLE serum (Fig. 2F). To better understand the effects of the serum on the organoids we also checked several other cytokines. We could observe significantly elevated levels of IL-18 and the presence of IL-6 in almost all SLE, but not control serum samples, as it has been reported for bigger SLE cohorts^41–45^ (Fig. 2G). Even though some of the control sera also showed increased levels of single cytokines only the SLE serum samples showed an overall increase of several cytokines (Fig. 2H). This result did not only indicate the overall inflammatory signature in SLE serum samples but also showed the diversity of the applied samples.

Taken together, our data revealed that organoids stimulated with serum derived from SLE patients were characterized by type I IFN signature similar to that of patient derived immune cells. The activation of IFN signaling underlined the so far neglected impact of IFN on epithelial cells in the context of SLE.

### 3. SLE serum stimulation increases mitochondrial respiration and leads to reduced fitness of organoids

We wanted to understand which other effects SLE serum would have on epithelial cells. Our focus was to investigate the influence of SLE stimulation on mitochondria, as these organelles are implicated in SLE pathogenesis^46, 47^ and serve as crucial regulators in maintaining intestinal homeostasis^48, 49^. The term mitochondrion represented 10% of all DEGs (Fig. 3A) while the enrichment analysis showed an upregulation of genes relevant for oxidative phosphorylation (Fig. 1F). Closer analysis revealed that mitochondrial encoded genes belonging to complex I and IV exhibited high expression levels in general (Fig.3A, B; Suppl. Fig.3A). Thus, their further upregulation in SLE stimulated organoids indicated their potential impact on respiratory chain activity. Evaluation of expression levels for each SLE serum donor demonstrated that this effect was induced by almost all samples (Fig. 3B and Suppl. Fig. 3A). To exclude that the overall upregulation of mitochondrial related genes was caused by an increase in mitochondrial mass we assessed the mitochondrial content in the serum stimulated organoids. Since it is known that the mitochondrial morphology is cell type dependent^50^ we decided to use flow cytometric analysis to quantify overall changes in mitochondrial mass. The results revealed that there were no differences in mitochondrial mass between both experimental conditions (Fig. 3C and Suppl. Fig. 3B). This indicated that the upregulated mitochondrial genes were due to an increased mitochondrial activity rather than caused by an increase of mitochondrial mass.

**Figure 3:**
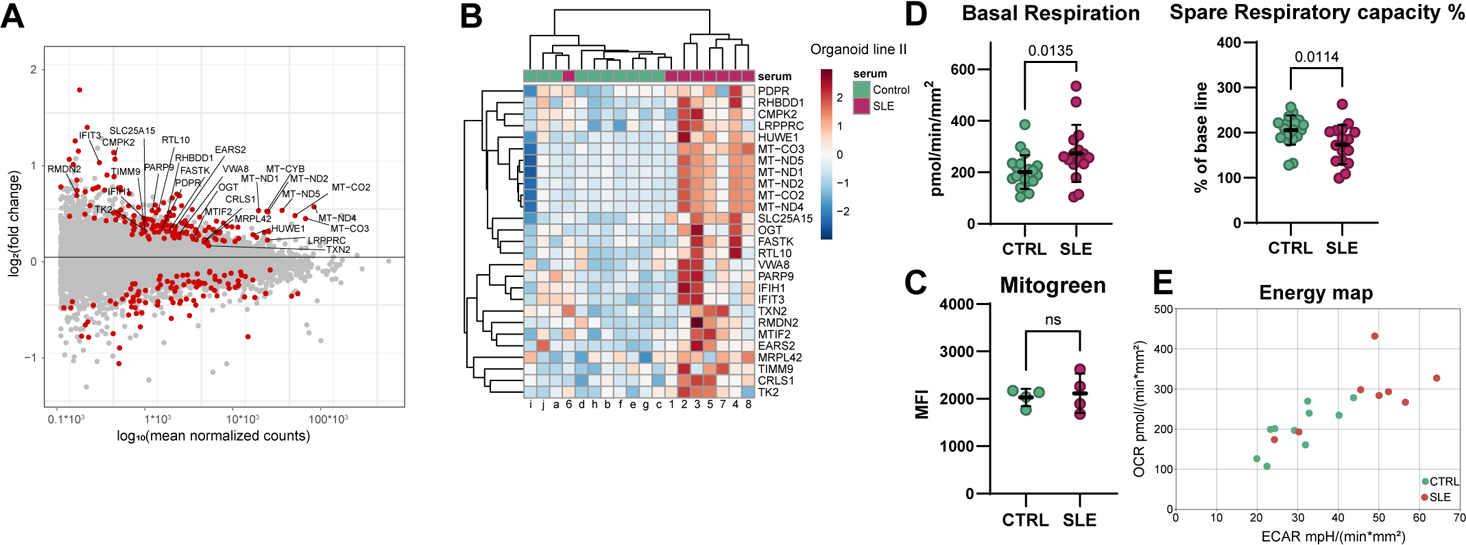
Effect of SLE serum stimulation on mitochondrial activity and organoid fitness. A: MA plot representing differential expression of all genes (DEGs) associated with the term Mitochondrion, indicating a significant proportion of upregulated DEGs to be mitochondria related. B: Heatmap demonstrating upregulation of mitochondrial genes in organoids stimulated with SLE serum, based on expression levels across various serum donors on organoid line II; n= 10 control serum (samples a to j) and n=8 SLE serum (samples 1 to 8). C: Mitogreen staining of organoids subjected to flow cytometry analysis, showing no significant difference in mitochondrial mass between control and SLE serum-stimulated organoids. Each dot represents one well of organoids dissociated into single cells and analyzed for mean fluorescence intensity (MFI); n=4 for each condition. D: Seahorse assay revealing increased basal respiration and diminished relative spare respiratory capacity, in SLE serum stimulated organoids. Each dot corresponds to one well of organoids stimulated with n=10 control serum samples or n=8 SLE serum samples. Each serum stimulation was analyzed with two technical replicates. E: Energy map demonstrating altered metabolic profile of organoids following SLE serum stimulation, with a shift toward higher glycolytic activity. Each dot corresponds to the average of two technical replicates of stimulated organoids, n= 10 control serum and n=8 SLE serum. Statistical tests performed with unpaired t-test.

We assessed mitochondrial function by a respiratory assay and saw a significant increase of basal respiration and ATP production along with an unaltered maximal respiration upon SLE serum stimulation (Fig. 3D and Suppl. Fig. 3C). Similar results were seen when CD8^+^ T cells were stimulated with IFNα.^46^ Upon assessing the relative spare respiratory capacity, a notable decrease was observed following SLE serum stimulation, which aligned with previous reports on altered mitochondrial function in CD8^+^ T cells derived from SLE patients.^46^ These results indicated that we were looking at an increased energy demand and a diminished capacity of intestinal epithelial cells to adjust to a dynamic energy demand after SLE serum stimulation.

Overall, we could see that the metabolic profile of the organoids was altered upon SLE serum stimulation (Fig. 3E). We were interested in understanding if this alteration was caused by a changed cell type composition of the organoids. Reports from literature suggest that more differentiated cells switch from glycolysis to oxidative phosphorylation as their primary energy source which would be reflected by a shift from the right lower to the left upper quadrant in the energy map.^50–53^ However, there is limited knowledge about the metabolic profile of different cell types in human colon. We therefore generated organoids with different cell type composition and observed a metabolic shift with a trend towards higher oxidative phosphorylation upon differentiation (Suppl. Fig. 3D and E). The metabolic shift we observed upon SLE serum differed and showed a trend to higher glycolytic activity compared to the control (Fig. 3E). While the impact of differentiation primarily relied on OCR (oxygen consumption rate), the effects of SLE stimulation influenced both OCR and ECAR (extracellular acidification rate). These results suggested that the changes induced by SLE serum stimulation had a more complex cause than just a shift towards altered differentiated phenotype and might be additionally caused by an overall change in mitochondrial function.

### 4. Expression of secretory cell markers are reduced while proliferation seems to be unaffected

Our *in vitro* model offered a unique opportunity to examine whether serum stimulation could affect differentiation dynamics, thereby replicating the potential impact on the continuous regeneration of the epithelial cell layer *in vivo* during SLE onset and progression. Cytokines have the capacity to shape the proliferation and differentiation dynamics.^54, 55^ However, not much is known about how epithelial cells specifically are influenced by cytokines at *in vivo* relevant concentrations contained in serum of SLE patients. Analysis of the proliferative capacity of organoids stimulated with SLE serum did not show any changes when measured by size, cell cycle (KI67), or S phase (EdU) (Fig. 1B, Suppl. Fig. 4 A and B). Since the hyperenrichment analysis revealed the terms secretory vesicle and secretory granule amongst the top 7 down regulated gene sets (Fig. 1G, 4B and Suppl. Fig. 4C) indicating an effect on the secretory function of the epithelial barrier we focused on cell type composition changes in the organoids. We checked a panel of cell type marker genes which showed almost exclusively effects on differentiated cells as indicated by a decrease of absorptive and secretory lineage markers (Fig. 4A). Of note, the absorptive cell markers SERPINA1, Serpin Family A Member 1, (also known as AAT) and SCNN1A, Sodium Channel Epithelial 1 Subunit Alpha, (also known as ENaC) are both connected to mucus layer build-up and function. SERPINA1 has antimicrobial functions^56^ while SCNN1A is important for ion and fluid regulation^57^ thereby playing a role in modifying mucus characteristics. Amongst the secretory cell markers were well known goblet cell markers AGR2, TFF3 and SPINK4 as well as CHGA which marks enteroendocrine cells. AGR2 is essential for the production and processing of gel-forming mucins such as MUC2^58^, whereas TFF3 forms a complex with FCGBP, one of the main components of mucus^59, 60^. We were therefore especially interested to understand if we could detect changes in the secretory cells. The number of goblet cells in stained sections was considerable variable between individual organoids, posing challenges for quantification. Analysis of MUC2 staining showed a significant decrease in intensity indicating an alteration in secreted mucus or changes in goblet cell function (Fig. 4C and D). The quantity of enteroendocrine cells, another type of secretory cell, decreased significantly, as did the amount of CHGA per enteroendocrine cell. (Fig. 4E, F and G). Overall, these results indicated that SLE serum stimulation alters the differentiation dynamics towards the secretory lineage and suggested a change in mucus composition marked by a reduction in MUC2, the major component of mucus, as well as other factors secreted to it.

**Figure 4:**
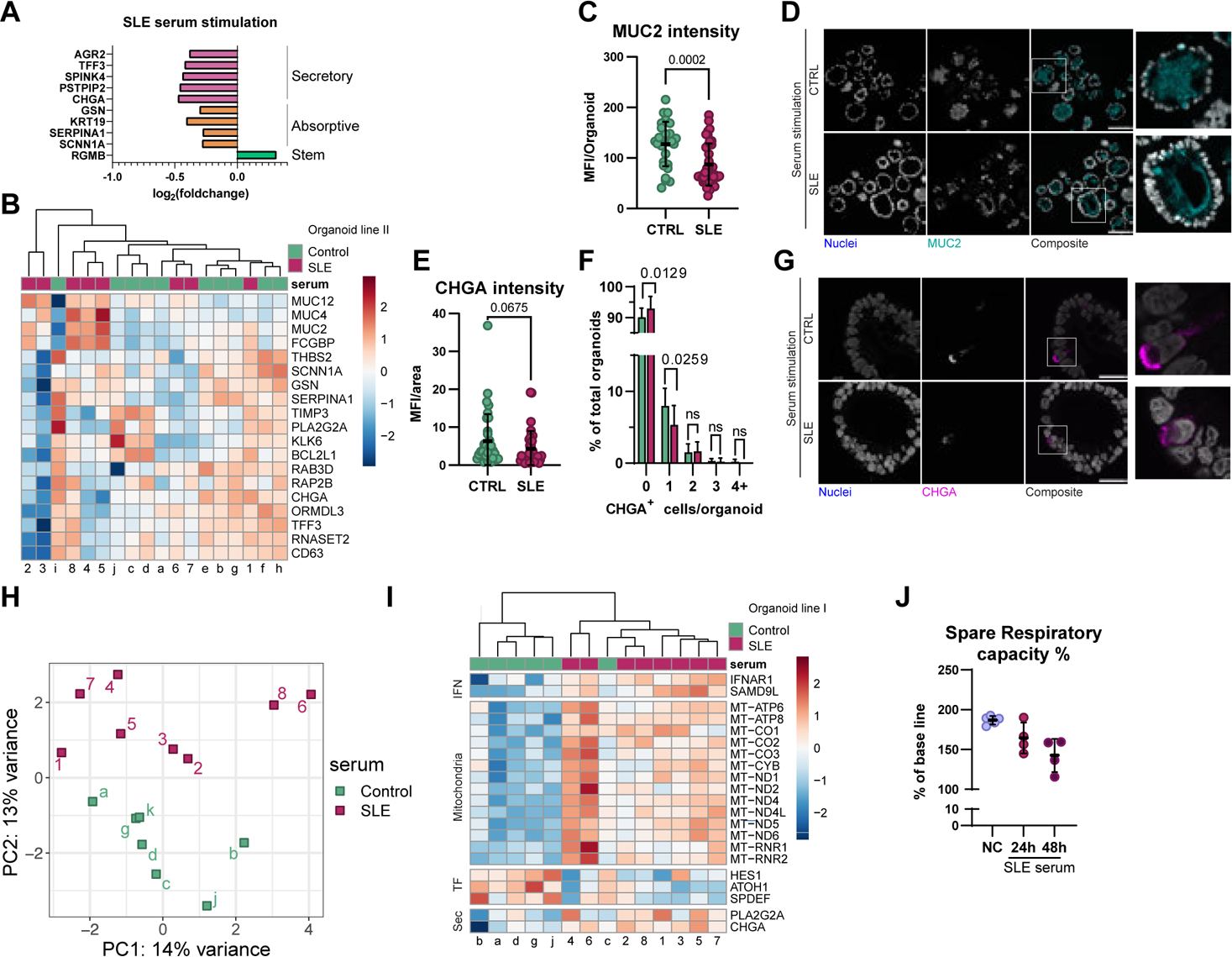
SLE serum stimulation impacts differentiation dynamics, particularly affecting the secretory lineage. A: Comparison of cell type marker expression levels in organoids stimulated with SLE and CTRL serum for 48h, highlighting a reduction in differentiated cells, as evidenced by the decrease in markers associated with absorptive and secretory lineage. B: Heatmap illustrating downregulated genes associated with the term secretory vesicle and mucus in 72h SLE serum stimulated organoids in organoid line II; n= 10 control serum and n=8 SLE serum. C: Quantification of mucus mean intensity per organoid normalized to the number of nuclei. Each dot corresponds to a single organoid, revealing a decrease in mucus intensity. Control n=29 organoids and SLE=40 organoids analyzed using unpaired t-test. D: Fluorescence microscopy images of organoids after 72h stimulation with either control serum (top) or SLE serum (bottom) associated with MUC2 staining analysis. Nuclei marked by DAPI and mucus by MUC2. Insets showcase enlarged areas, scale bars 100 μm. E: Quantification of CHGA intensity normalized by the area of the cell determined by DIC (Differential Interference Contrast) microscopy. Each dot represents a single organoid showing a decreased CHGA abundance. Control n=36 organoids and SLE=34 organoids (analyzed using unpaired t-test). F: Distribution of number of CHGA positive cells per organoid normalized to total organoid number (analyzed by ANOVA with Holm-Šidák’s multiple comparisons test) suggesting a decrease in enteroendocrine cells. n=1270 organoids for control and n=801 organoids for SLE serum stimulation were analyzed. G: Confocal microscopy images of organoids after 72h stimulation with either control serum (top) or SLE serum (bottom) associated with CHGA staining analysis. Nuclei marked by DAPI and enteroendocrine cells by CHGA. Insets show enlarged areas, Scale bars 25 μm. H: PCA plot illustrating distinct clustering following 24h stimulation with SLE or control serum, indicating an early impact at transcriptomic level. I: Heatmap representing upregulated genes connected to IFN signaling, mitochondria, transcription factors related to differentiation (TF) and secretory lineage (sec) after 24h stimulation, indicating their role in initiating the observed changes. J: Seahorse assay revealed a gradual decrease in relative spare respiratory capacity in organoids stimulated with SLE serum for 24 and 48h compared to unstimulated controls (NC). This decrease is similar to what was observed after 72h of stimulation. Each dot on the graph represents a single well of organoids; n=5 for NC and n=4 for control and SLE serum stimulation. Unless otherwise specified, all experiments depicted in this figure were analyzed with n= 10 control serum (samples a to j) and n=8 SLE serum (samples 1 to 8).

### 5. Secretory lineage differentiation is affected 24h after stimulation

To better understand the underlying mechanism, we stimulated organoids for 24h with SLE or control serum. This gave us the opportunity to confirm that we were facing an effect on the differentiation process rather than cell type loss. Sequencing analysis indicated that even brief stimulation could trigger a distinct response, evidenced by the separate clustering in the PCA analysis (Fig. 4H). Amongst the DEGs we observed an upregulation of genes connected to IFN signaling and several mitochondrial encoded genes indicating their distinct role in initiating the SLE serum-induced changes (Fig. 4I). Functional analysis showed a stimulation time-dependent decrease of relative spare respiratory capacity similar to that observed after 72 h stimulation (Fig. 4J and Fig. 3D). Additionally, transcription factors, SPDEF, ATOH1 and HES1, all important for the induction of differentiation, were down regulated indicating a delayed or altered differentiation towards specialized cells (Fig. 4I).

In summary, these results support the hypothesis that SLE serum stimulation induced alterations in differentiation, particularly impacting the secretory lineage. Short-term serum stimulation induced changes in transcription factor expression necessary for differentiation which upon long-term stimulation leads to alterations in cell type composition of the organoids. Ultimately, the consequence will accumulate in an altered mucus layer that *in vivo* has the potential to trigger alterations in the protective function of the mucus and influence the microbiome.

### 6. Effects on epithelial cells depend on IFNAR1, although not being exclusively linked to IFNα activity

The importance of type I IFN in SLE pathogenesis and its potential as target for treatment was highlighted by the approval of anifrolumab, a type 1 interferon receptor (IFNAR1) antagonist in 2021.^61^ Our novel disease model was able to show the involvement of type I IFN in altering epithelial cell signatures as suggested by the increased IFNSG expression and activation of IFN signaling cascade (Fig. 2A-E). To verify this, we utilized the inhibitory potential of anifrolumab on IFN signaling. We validated that anifrolumab was able to inhibit phosphorylation of STAT1, a downstream target of IFNAR1 activation (Suppl. Fig. 5A) by organoid stimulation with up to 100 ng/ml IFNα2b. Analysis of the expression profile showed no significant difference of organoids treated with anifrolumab in addition to serum comparing SLE to CTRL serum stimulation (Fig. 5A). To further validate the successful inhibition of IFNAR1 signaling we checked normalized counts of DEGs specific for epithelial IFN signaling identified before (Fig. 5B and 2B). We could see that anifrolumab was able to abolish the increased expression of these genes not only in IFN stimulated organoids but more importantly also in combination with SLE serum stimulation making them indistinguishable to the control condition (Fig. 5B). As a next step we wanted to assess if anifrolumab was also able to revert the mitochondrial dysfunction. The respiratory assay confirmed that with blocking IFNAR1 SLE serum stimulated organoids had a similar relative spare respiratory capacity as in the control condition (Fig. 5C). Additionally, to confirm that the effect we were looking at was highly specific to SLE, we analysed the effect of serum from patients suffering from another systemic autoimmune disease, granulomatosis with polyangiitis (GPA). We could show that organoids stimulated with serum from GPA patients, a disease not driven by type I interferon, showed no distinctive expression pattern (data not shown) and no mitochondrial dysfunction underlining the dependency on type I interferon (Suppl. Fig. 5B).

**Figure 5:**
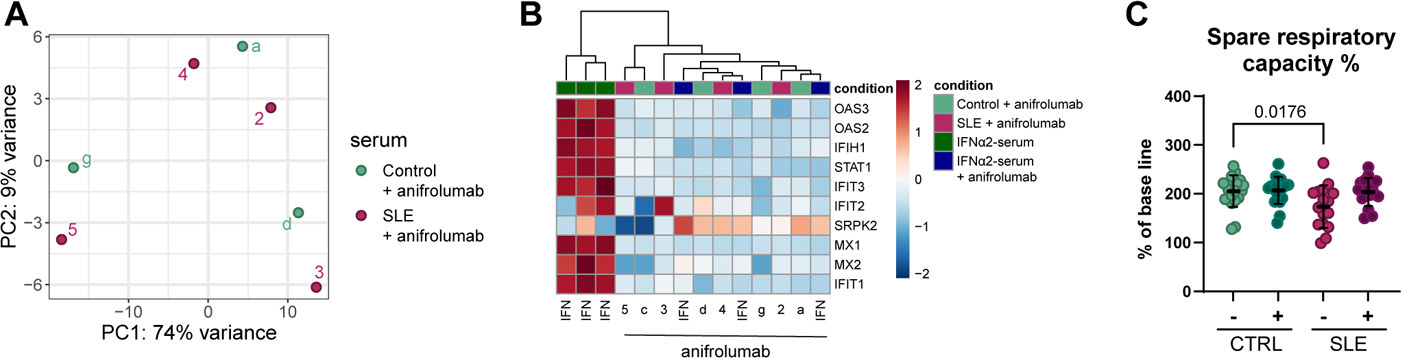
Visualization the effects of blocking IFNAR1 with anifrolumab on organoid stimulated with either SLE or CTRL serum. A. PCA plot illustrating the gene expression profiles of organoids treated with anifrolumab in addition to SLE or CTRL serum stimulation. The plot shows no significant difference in expression profiles between these treatment groups, emphasizing the role of type I interferon and inhibition of IFNAR1 signaling. n= 4 control serum + anifrolumab (samples a, c, d and g) and n=4 SLE serum (samples 2 to 5). B. Heatmap showing differentially expressed genes (DEGs) specific to epithelial IFN signaling before and after treatment with anifrolumab. Noticeable reduction in the expression of these genes is evident following anifrolumab treatment, confirming successful inhibition of IFNAR1 signaling as seen in the positive control (IFN). n= 4 control serum + anifrolumab (samples a, c, d and g) and n=4 SLE serum (samples 2 to 5), n=3 IFNα+control serum, and n=3 IFNα+control serum+anifrolumab. C. Results from Seahorse assay measuring relative spare respiratory capacity in organoids. Blocking IFNAR1 with anifrolumab resulted rescue of the phenotype induced by SLE serum stimulation, suggesting the mitigation of mitochondrial dysfunction. Statistical test performed with one way ANOVA with Holm-Šidák’s multiple comparisons test. Each dot corresponds to one well of organoids stimulated with n=10 control serum samples or n=8 SLE serum samples. Each serum stimulation was analyzed with two technical replicates.

To confirm that the effect we observed was specific to SLE serum, rather than simply a consequence of IFN independently, we stimulated organoids with IFNα in combination with control serum (IFNα-serum). However, this stimulus was not able to induce the same complex response as seen for SLE serum. Overlapping the DEGs from both comparisons, SLE compared to control serum and IFNα-serum compared to IFNα-serum+anifrolumab, showed only a small overlap of almost exclusively IFN related genes highlighting the unique response of the organoids to SLE serum (Suppl. Fig. 5C).

Collectively, these findings suggest that the response of the organoids to SLE serum is dependent on IFNAR1, but not caused solely by its activation by IFNα. This implies that the unique composition present in the SLE serum plays a significant role in provoking the response, in conjunction with the dependency on IFNAR1.

### 7. Colon organoid Single cell sequencing reveals the presence of all major colonic cell types found *in vivo*

To gain a deeper understanding of the cell types that could potentially exhibit higher sensitivity to serum stimulation, as well as to examine the cellular composition of the organoids, we conducted single-cell RNA sequencing. This approach allowed us to analyze expression patterns of individual cells and gain insights into their specific responses to serum stimulation. We could assign nine clusters to the cells analyzed which showed a distinct cell cycle distribution and expression profile specific to their cell type (Fig. 6 A-C; Suppl. Fig. 6A and B). Trajectory analyses identified lineage transitions and showed connectivity between progenitor and differentiated populations (Fig. 6D and Suppl. Fig. 6C). As anticipated, stem cells gave rise to transit-amplifying cells (TA1, TA2 and TA3), the secretory lineage (GC) and the absorptive lineage comprising three colonocyte clusters (eCL, cCL and ncCL).

**Figure 6:**
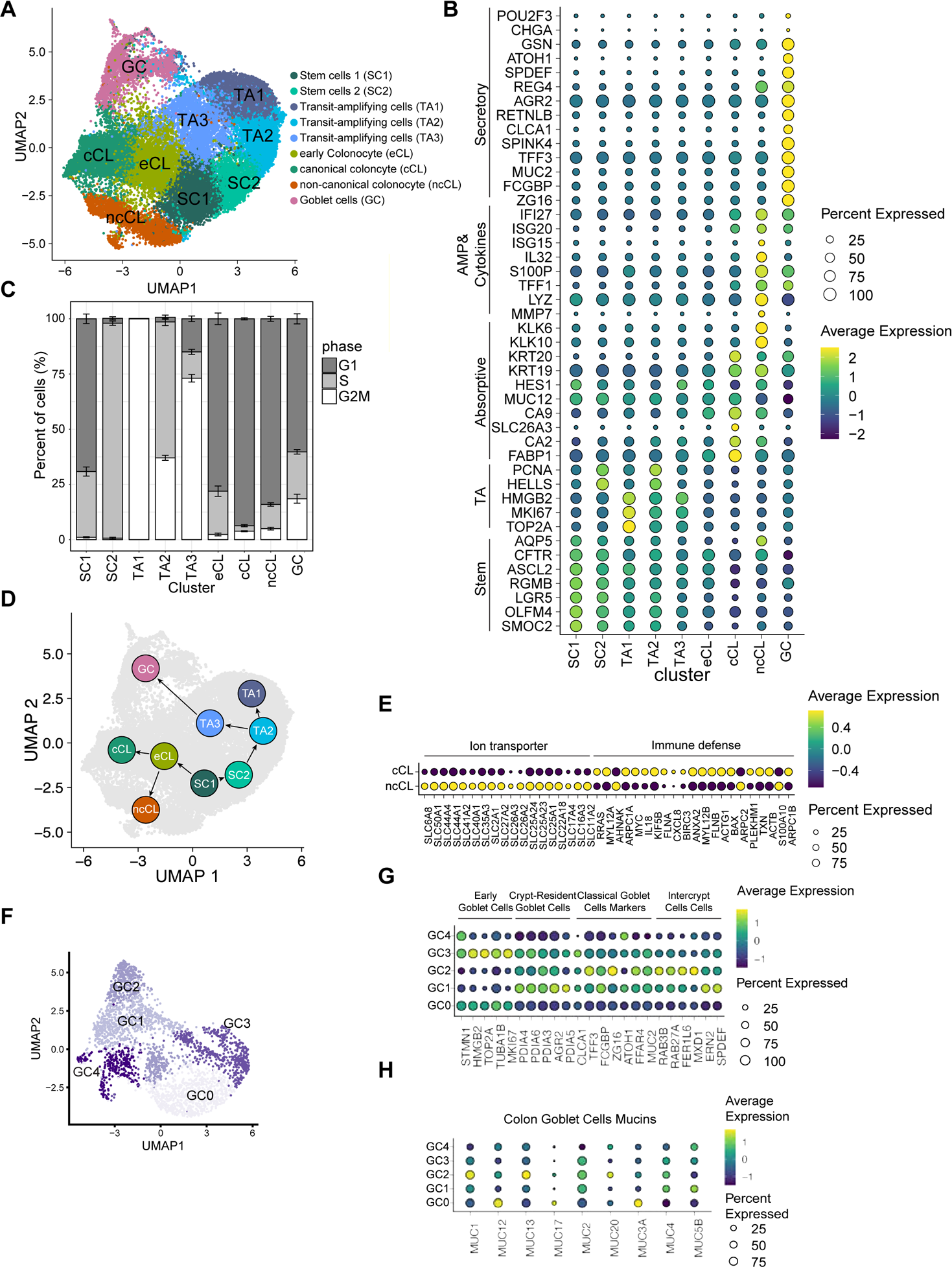
Comprehensive single-cell RNA sequencing reveals diverse cellular composition in colon-derived organoids. A. UMAP demonstrates the distribution of all examined cells derived from single-cell RNA-Seq data. These cells come from control (n=3) and SLE (n=3) serum-stimulated organoids and are color-coded to represent 9 distinctive clusters. B. Dot plot illustrating the predominant marker genes corresponding to each cluster in colon-derived organoids. The size of the dots corresponds to the proportion of cells expressing a specific gene, while the colour indicates the mean relative gene expression across the respective cell type. C. Cell-cycle phase distribution for each cluster, delineated by cell type. Mean with SEM is shown. D. Diagram representing the summary of the different predicted cellular developmental trajectory using Slingshot. E. Dot plot of canonical and non-canonical colonocyte marker genes of the identified absorptive colonocyte clusters cCL and ncCL. Dot size corresponds to the percentage of cells expressing each gene, while the colour indicates the mean relative gene expression across the cell type. F: UMAP representation showing subclustering of goblet cells and variation in cell populations. G: Dot plot representing the average expression of goblet cell subtype marker genes across identified goblet cell subclusters. Dot color and size indicate average gene expression and the proportion of cells expressing the gene, respectively. H: Dot plot displaying the unique combinations and average expression levels of mucin genes across goblet cell subclusters. Dot color and size represent average gene expression and the percentage of cells expressing the gene, respectively.

We identified two distinct clusters of stem cells (SC1 and SC2) marked by the expression of classical stem cell markers LGR5, OLFM4, RGMB, and SMOC2 (Fig. 6A, B). The key characteristic that sets apart these two clusters is their distinctive cell cycle distribution. While the majority of cells in SC2 are in S phase, a significant portion of cells in SC1 are in G1 phase (Fig. 6C). Transition to cycling TA cells (TA1) was marked by the transition to exclusively G2/M phase and an enrichment of the proliferation markers MKI67 and TOP2A expression (Fig. 6B and C). Furthermore, we identified two additional TA clusters, TA2 and TA3 (Fig. 6A). All three TA clusters showed gene-expression gradients along with distinct cell cycle distributions reflecting active proliferation and transition to progenitors of absorptive and secretory lineage (Fig. 6B and C; Suppl. Fig. 6C). Cell transition to G1 phase marked all identified differentiated cell types, early colonocytes (eCL), canonical colonocytes (cCL), non-canonical colonocytes (ncCL) and goblet cells (GC) (Fig. 6C).

Interestingly, we could identify two different types of colonocytes emerging from early colonocytes (Fig. 6D). Canonical colonocytes were marked by the expression of classical absorptive markers SLC26A3, CA2 and FABP1. These markers were absent in the non-canonical colonocytes which were marked by high expression of MMP7, LYZ, IL32 and the IFN response genes ISG15, ISG20 and IFI27 (Fig. 6B). Even though MMP7 and LYZ were lately described to be present in deep crypt secretory cells^62, 63^, our cell population lacked other reported markers of these newly described cells. In addition, we could observe the expression of HES1 an exclusive marker of the absorptive lineage (Fig. 6B).

Since the clustering showed two distinct populations, we wanted to understand the major features separating them. Analysis of the significantly enriched markers (p_adj_<0.1 and log_2_fc >0.25) suggested different functions of the two cell types. Canonical colonocyte showed an enrichment of lipid metabolism related pathways and expressed several ion transporters. Specifically SLC26A3 and CA2 which *in vivo* are expressed by fully mature colonocytes, the central players of absorption^13^ suggesting that also *in vitro* their function would focus on lipid processing and absorption. In contrast, we could observe an enrichment of pathways connected to immune defense in non-canonical colonocytes indicating cytokine-driven responses to maintain barrier integrity (Fig. 6E).

The secretory lineage was characterized by the presence of classical goblet cell markers MUC2, SPINK4, SPDEF and ATOH1. In addition, we could see the expression of ZG16, FCGBP and CLCA1, all mucus components (Fig. 6B). Further subclustering of the goblet cell population resulted in 5 clusters (Fig. 6F and Suppl. Fig.6D). These could be assigned to two early goblet cell types (GC0 and GC3) and two types of differentiated goblet cell populations (GC1 and GC2) (Fig.6G; Suppl. Fig. 6E). Analyzing their gene expression pattern revealed that cluster GC1 had increased expression of AGR2, FCGBP, CLCA1, ERN2 and SPDEF all known to be mucus core proteins or genes involved in mucus biosynthesis^64^(Fig. 6G). In contrast to that cluster GC2 was characterized by a higher expression of ZG16 and TFF3 which function as AMP and regulator of mucus viscosity respectively^65, 66^. Cluster GC4 showed only low expression levels of typical goblet cell markers. With further analysis we were able to identify a small fraction of cells within this GC4 expressing CHGA and POU2F3 indicating the presence of EECs and Tuft cells respectively (Suppl. Fig 6F). Interestingly, similar as reported previously for human colon tissue^67–69^ we were able to see differences in mucin gene expression patterns between the goblet cell cluster indicating their distinct functionality also *in vitro* (Fig. 6H).

In summary, we identified nine unique clusters composed of stem cells, TA cells, goblet cells, canonical and non-canonical colonocytes. Additionally, we observed 5 subclusters within the secretory cell population including enteroendocrine and Tuft cells as well as different goblet cell subtypes. Thus, our *in vitro* model represents the majority of the cell types found in descending colon providing a valuable model to study the effect of SLE serum on an *in vivo* like intestinal epithelial barrier.

### 8. scRNA-seq analysis unveils diverse cellular responses leading to an alteration in mucus composition

After confirming the presence of the major colonic cell types in our near-physiological *in vitro* model we wanted to understand which impact the SLE specific serum signature would have on different epithelial cell types. Differential gene expression analysis within all identified cells revealed 480 genes (with a p_adj_ <0.1 and average log_2_fold change >|0.1|) that were misregulated upon SLE serum stimulation. We observed downregulation of pathways related to protein translation and altered expression of genes connected to oxidative phosphorylation specifically in stem cells, early colonocytes and goblet cells (Suppl. Fig. 7A and B). Both protein translation and mitochondrial function are important players in modulating proliferation and differentiation underlining the altered cell type composition upon SLE serum stimulation.^70, 71^ Further analysis showed that some genes were broadly altered like Macrophage Migration Inhibitory Factor (MIF),the tight junction protein Claudin 3 (CLDN3) or AQP5 which were significantly downregulated (Fig.7A). However, we could see that each cell type had a specific response to SLE serum stimulation. In SC1, but not in SC2, stem cell markers SMOC2 and OLFM4 were significantly downregulated. In contrast, in canonical colonocytes typical absorptive markers were unaltered while tight junction proteins TJP1 and CLDN7 were downregulated. Both are important for barrier integrity and their absence is associated with a delay in mucosal repair or an initiation of inflammation *in vivo*.^72, 73^ This specifically suggests an alteration in the epithelial barrier and lays the ground for a self-perpetuating chronic inflammation a seen in SLE.

**Figure 7:**
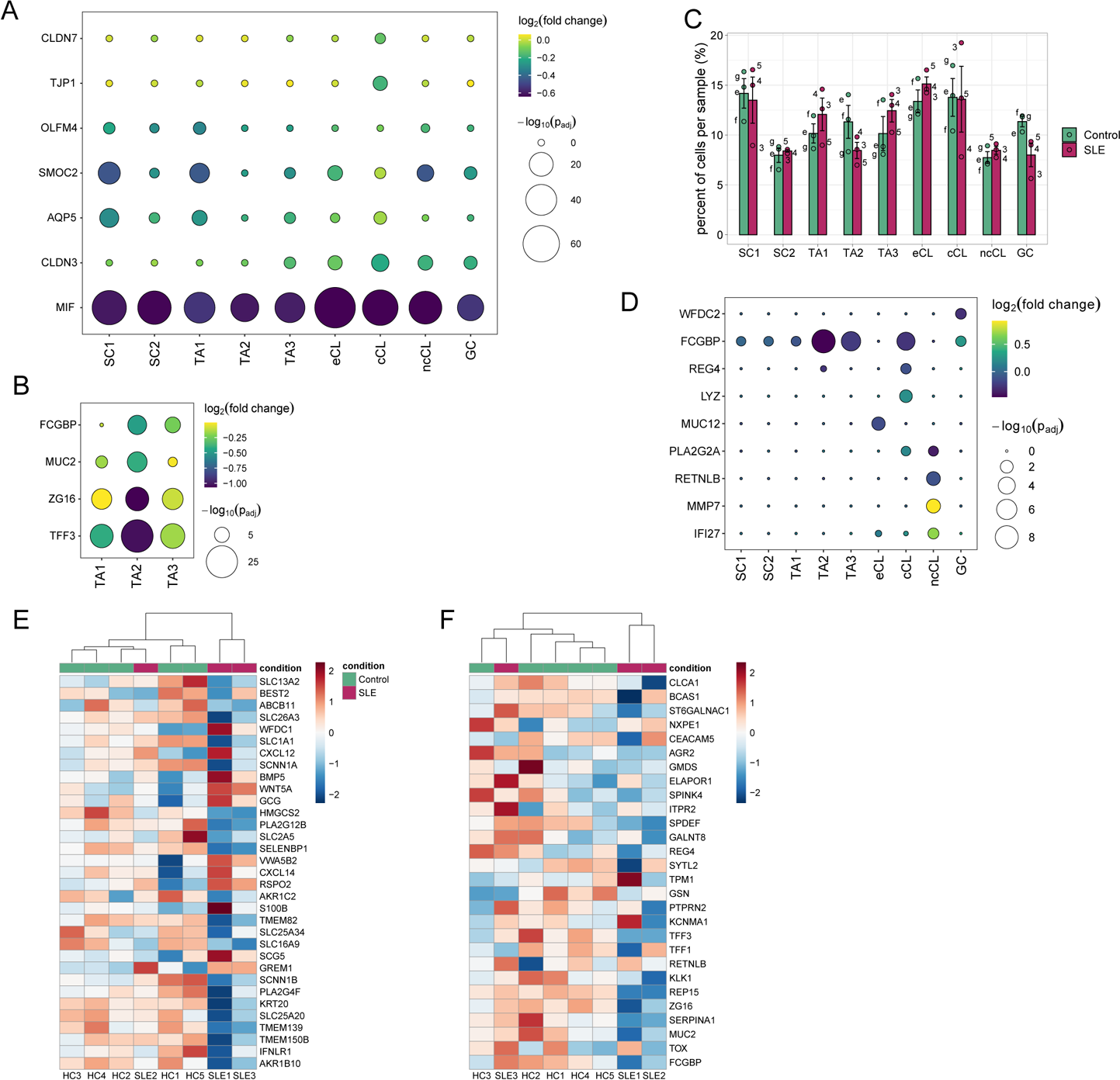
Impact of SLE serum stimulation on organoid cellular composition and goblet cell diversity. A: Dot plot depicting differential expression after SLE serum stimulation. This figure illustrates the cell cluster specific transcriptional alterations caused by SLE serum, with downregulated genes such as MIF, CLDN3 and AQP5 across multiple cell types, and distinctive responses like decreased SMOC2 and OLFM4 in stem cells and decrease of TJP1 and CLDN7 in canonical colonocytes. B: Dot plot demonstrating the impact of SLE serum on the expression of specific goblet cell markers in TA (TA1, TA2, TA3) cells. C: Bar graph displaying the percentage of cell populations in each cluster when comparing control samples with those stimulated by SLE serum. n= 3 control serum (samples e to g) and n=3 SLE serum (samples 3 to 5). D: Dot Plot of representing the fold change of gene expression alterations in absorptive and goblet cells following SLE serum stimulation compared to controls. The size and color of each dot represent p-adjusted value and fold change, respectively. H: Heatmap representing differential gene expression of descending colon biopsies from healthy controls (n=5) and SLE patients (n=3). I: Heatmap representing differential expression of genes from human descending colon biopsies. Shown are genes from a curated list of goblet cell markers based on findings *in vitro*. healthy controls (n=5) and SLE patients (n=3).

Next, we wanted to understand if we could detect a change in goblet cell differentiation as indicated by the bulk RNA sequencing data. We observed a noticeable decline in the levels of goblet cell markers such as TFF3, ZG16, MUC2, and FCGBP within the TA population (TA1, TA2, and TA3) (Fig. 7B). This suggested that the stimulation with SLE serum led to diminished differentiation towards the secretory lineage, which in turn suggested a potential reduction of cells differentiating towards goblet cells. Our findings were corroborated by the results from the bulk RNA sequencing analysis, wherein we noticed a reduction in the expression of early goblet cell markers during the initial phase of stimulation (24h), and a subsequent decrease in late secretory markers with extended stimulation (72h) (Fig. 4A, B, I). Consistent with these results, we could detect a minor reduction in the number of cells in the goblet cell cluster (Fig. 7C and Suppl. Fig. 6A). Interestingly, within the diminished goblet cell population we observed a significant upregulation of FCGBP, an IgGFc-binding protein, which is crucial for structural integrity of the mucus layer^60, 74^. Additionally, WFDC2 which was only recently identified to be required for barrier integrity and is downregulated in goblet cells of UC patients^75^ was also downregulated upon SLE serum stimulation (Fig. 7D). Further analysis showed generalized trend towards a decrease across all goblet cell subclusters, with a significant reduction in GC4 (Suppl. Fig. 7C). GC4 lacked the expression of classical goblet cell markers and contained CHGA expressing cells suggesting a reduction in enteroendocrine cells as identified by staining (Fig. 4F). Additionally, GC4 showed no DEGs while the DEGs of the other GC clusters were mainly downregulated ribosomal genes connected to translation and protein translocation. This might indicate an impact on goblet cells function that highly depends on a constant translation and protein trafficking. Apart from the reduced number of goblet and enteroendocrine cells, these findings suggest an altered mucus synthesis.

Since mucus properties can easily be modified by an altered abundance of mucins and AMPs^76, 77^ which is not restricted to goblet cells we checked known markers in the absorptive lineage. In the early colonocyte population we observed a downregulation of MUC12 a membrane bound mucin which is part of the protective glycocalyx^12^. In addition, we observed an increase of lysozyme (LYZ) and decrease in REG4 in the canonical colonocytes (Fig. 7D). Both are known to be secreted into mucus and are important for microbial defense^78, 79^. A similar alteration was also evident within the non-canonical colonocytes where we observed an alteration in the expression of MMP7, RETNLB and PLA2G2A, genes that are essential components of barrier function, especially RETNLB and PLA2G2A with their bactericidal properties^80–83^ (Fig. 7D). Additionally, in non-canonical colonocytes and GC2 we were able to detect the induction of IFI27, an interferon inducible gene, similar as reported for whole blood samples from SLE patients ^84–86^ (Fig. 7D) and as observed by the induction of IFNSG in our bulk RNA sequencing data (Fig. 2C). Interestingly, non-canonical colonocytes and GC2 both expressed AMPs which are known to be altered by proinflammatory cytokines^87–89^ suggesting a similar connection in our model.

Overall, these results revealed that SLE serum stimulation of epithelial cells lead to distinct transcriptional alterations which are marked by a shift in differentiation as already suggested by our bulk RNA sequencing data. The observed expression changes in goblet cell markers, mucins and AMPs in both, the secretory and absorptive lineage, along with the reduction of goblet cell number might integrate in an ultimately changed mucus composition and compromise its function as first line of defense. *In vivo* such changes could have detrimental effects on the barrier integrity, immune response, and the gut microbiome

### 9. Intestinal epithelial barrier is altered in descending colon tissue biopsies from SLE patients

To highlight the relevance of this disease model and provide context regarding intestinal involvement in SLE, we conducted an expression analysis of intestinal biopsies derived from descending colon of healthy controls (n=5) and SLE patients (n=3). We could identify 256 DEGs which were representing the integrated response of all present cell types in the biopsies. In our further analysis we focused on genes that were expressed by epithelial cells. Overall, we could observe a downregulation of several genes connected to absorption and ion transport (SLC1A1, SLC13A2, SLC16A9, SLC25A20, SLC25A34). BEST2, a marker for goblet cells, was significantly downregulated while markers for enteroendocrine cells (SGC5, GCG, VWA5B2) were upregulated suggesting an alteration in the secretory lineage (Fig. 7E). Additionally, we observed reduced expression of ion transporters SLC26A3, SCNN1A and SCNN1B while other markers for colonocytes were unaltered. This indicated an alteration in cell function rather than a loss of colonocytes. Those three ion transporters are the major players in water absorption in descending colon and are able to modulate mucus viscosity, ion content and ultimately alter its structural composition via changes in sodium absorption and bicarbonate secretion influencing the mucosal pH.^13, 90^ In addition to that several factors important in the maintenance of the intestinal stem cell niche and differentiation (WNT5A, RSPO2, GREM1, BMP5) were significantly downregulated suggesting changes in the physiological proliferation and differentiation dynamics. We then checked the expression of the top secretory cell markers identified with single cell sequencing. Surprisingly, we could see clustering of two SLE patient samples that showed a reduction in the selected markers (Fig. 7F).

The evidence we have gathered collectively suggests that the impact of SLE is not confined to immune cells, but also affects intestinal epithelial cells. Specifically, we have observed significant alterations in the intestinal stem cell niche and the absorptive lineage. Changes in secretory cells echoed the results obtained from our *in vitro* model, further illustrating the multifaceted influence of SLE. Thus, these combined findings indicate that SLE pathogenesis involves epithelial cells and could potentially weaken the structural integrity of the mucus layer and disrupt the barrier function. This sheds light on the broader implications of SLE, revealing that its effects extend beyond the immune cell compartment.

## Methods

### 10. Human samples and study approval

Healthy and SLE whole human intestines were obtained from multiorgan donor patients that were rejected for transplantation due to reasons unrelated to intestinal diseases. Organoid line I and II were generated from a healthy 46-year-old female and a 23-year old male descending colon tissue respectively. Serum SLE and control samples were respectively obtained through collaborations with the Division of Rheumatology, Department of Medicine V, University Hospital Heidelberg, Germany and Molecular Neuroimmunology Group, Department of Neurology, University of Heidelberg, Germany. Further information about the serum samples can be found in Suppl. Table 1.

The protocol for serum collection was approved by the ethics committees of the University of Heidelberg (272/2006 and S-187/2008). Intestines that could not be used for transplant were procured from human donors via 1) Novabiosis, Inc. (Research Triangle Park, Durham, North Carolina, USA). The acquisition of the donor intestines was conducted following ethical committee approval from the Organ Procurement Organizations (OPO), in line with the consent and deidentification guidelines established by the OPOs and the United Network for Organ Sharing (UNOS), under the US transplantation network framework. Permission for organ donation was granted by the immediate family members of the donors while preserving the patient’s privacy. This donation approval aligns with the guidelines provided by the federal organization UNOS and the Federal Drug Administration (FDA). 2) Samples and data from patients included in this study were provided also by the I3PT Biobank and the DTI Foundation (Barcelona, Spain) they were processed following standard operating procedures with the appropriate approval of the Ethics and Scientific Committees.

All research procedures were conducted adhering to the principles stipulated in the WMA Declaration of Helsinki. All participants in the study gave written informed consent before their inclusion and were assured that their identities would remain confidential in relation to this study.

### 11. Organoid generation and culture

Crypt isolation was performed as previously published^91^. The mucosa of approximately 1 cm^2^ was separated from the underlying tissue and cut into small pieces, washed with cold PBS before incubation in 10 mM EDTA/PBS for 1 h at 4°C under constant rocking. The biopsies were then transferred to 5 ml cold PBS and crypts were liberated by pipetting 20 times. This was repeated twice, and the pooled fractions were centrifuged at 250g for 5 min at 4°C. The pellet was once washed in base medium (Suppl. Table 2) before the crypts were resuspended in 45 µl 80% Matrigel (#356231, Corning) in expansion medium (Suppl. Table 2) and plated in small drops in a 24-well plate. After polymerization 500 µl growth medium (Suppl. Table 2) containing Y-27632 was added until the first passage or later for the first days after passaging. Initial passage was performed when organoids reached a size of approximately 200-300 µm diameter. After establishing the organoid line organoids were passaged at a ratio of 1:6-1:10 every 4-5 days by a 45s incubation in Accutase at 37°C followed by mechanical dissociation in 6 ml base medium with a 10 ml pipet equipped with a 200 µl pipet tip. Organoid fragments were centrifuged at 150g for 5 min at 4°C and plated as stated above. Medium was changed every 2-3 days.

### 12. Organoid stimulation

Organoids were passaged as described above with the modification of excluding bigger sized fragments by using a 70 µm strainer after the dissociation. In addition, fragments were counted and 400 fragments/µl were seeded. Organoids were grown in expansion medium for four days before dissociating them into single cells by incubation for 90 s in Accutase followed by mechanical dissociation as described above. Bigger fragments were removed by using a 40 µm strainer. Then 800 cells/µl were seeded in 10 µl Matrigel mix in 96-well plate. After polymerization 100 µl growth medium were added containing Y-27632. Organoids were stimulated with 5% serum or 5% control serum in stimulation medium (Suppl. Table 2). In case of IFNα stimulation 100 pg/ml IFN-α2B (Humankine, HZ-1072) were added additionally. Anifrolumab was added at a final concentration of 10 µg/ml. Stimulation started either 24 h, 48 h or 72 h before harvesting organoids on day five.

### 13. Organoid quantification

To analyze organoid morphology brightfield images of the total Matrigel drop area were acquired at three different z positions in a range of 300 µm with Nikon Eclipse Ti2 inverted microscope. Images were analyzed using OrgaQuant tool ^92^ with settings adjusted to 1600 (image size), 2 (contrast) and 0.7 (confidence threshold). Average diameter was calculated.

### 14. LDH cytotoxicity assay

Supernatant of stimulated organoids was collected after 48 h incubation, diluted 1:100 in storage buffer and analyzed as indicated in the manufacturer’s protocol (LDH-Glo Cytotoxicity Assay, Promega, #J2381).

### 15. Immunohistochemistry

Organoids from two wells were pooled in base medium. The organoid pellet was resuspended in 100 µl harvesting solution (Cultrex, Biotechne) and incubated for 45 min at 4°C. The released organoids were washed with ice-cold PBS, then 4% PFA was added, and organoids were fixed for 45 min at 4°C followed by a final wash. The organoids were then resuspended in 4% low-melt point agarose. Paraffin embedding was performed following standard protocols. For immunohistochemistry 3 µm sections were rehydrated, heat-induced antigen retrieval in 10 mM sodium citrate acid and 0.05% Tween 20 (pH 6) and quenching 50 mM NH_4_Cl was performed. Sections were then blocked with 5% goat serum for 1 h. Primary antibodies anti-cleaved Caspase 3 (1:100; Cell Signaling, 9661S), anti-KI67 (1:200; abcam, ab15580), anti-MUC2 (1:200; Santa Cruz Biotechnology, sc-515032) or anti-CHGA (1:100; Novus Biologicals, NB120) were incubated overnight at 4°C. Anti-rabbit-488 (1:500; Invitrogen, A11034) anti-mouse-568 (1:500; Invitrogen, A-11031) was incubated together with Hoechst-33342 (Invitrogen, H3570) for 1 h at RT. Mounted sections were imaged using Nikon Ti-2 fluorescence or AX confocal microscope. Image analysis was performed using ImageJ^93, 94^. Nuclei were quantified with StarDist^95^ and overlap with KI67 quantified using BioVoxxel tool box^96^.

### 16. Western blot

Organoids were released from the Matrigel as described above. For protein isolation organoids from 4 wells per condition were pooled. The organoids were lysed in 120 µl RIPA buffer (Thermo Scientific) containing 1.4x PhosphoStop and cOmplete mini (Roche) for 45 min on ice. Protein concentration was determined using BCA Protein Assay Kit (Pierce). Lysates were diluted in Laemmli buffer at a concentration of 0.5 µg/µl, 20 µl were loaded for SDS-PAGE and then transferred to a PVDF membrane. The membrane was blocked with 5% milk in TBST for 1 h at room temperature and then probed overnight at 4°C with pStat1 (1:1000; Cell Signaling #9167S), β-actin (1:500; Cell Signaling #4970S), STAT1 (1:1000; Cell Signaling #9167S) or IFIT3 (1:500; Santa Cruz #sc-393512). Signal was detected after incubation with anti-mouse_HRP (1:2000; Cell Signaling, 7076P2) or anti-rabbit-HRP (1:2000; Cell Signaling, 7074P2) for 1 h at room temperature using ECL solution Clarity Max (Bio-Rad Laboratories) and fluorescence-imaging device (Vilber, FUSION FX7).

### 17. IFNα/β receptor cell line

IFN-α/β reporter HEK-Blue Cells (InvivoGen, #hkb-ifnab) were selected and grown to confluency in a 96-well plate. Cells were stimulated for 24 h with 10% serum. The supernatant was then mixed with QuantiBlue Solution (Invivogen, #rep-qbs), incubated 3 h at 37°C and alkaline phosphatase (SEAP) activity determined measuring the absorbance at 600 nm using GloMax®-Multi Detection System (Promega).

### 18. Serum analysis

Serum samples were analyzed using a bead-based multiplex assay (BioLegend, LEGENDplex, 740809) following the manufacturer’s protocol. 5000 events (bead populations A + B) were recorded using a BD FACS Aria Fusion flow cytometer. Cytokine concentration was calculated based on a standard curve using Biolegend’s LegendPlex data analysis software. Cytokine levels that were below the detection limit were set to half the detection limit to calculate statistical significance. Cytokine levels for each sample can be found in Suppl. Table 6.

### 19. Cytometric analysis

Single cell suspensions were prepared, and cells stained with MitoTracker green (10 nM), 50 nM MitoTracker red (Invitrogen) for 15 min at 37°C and 5% CO_2_ atmosphere. The cells were then washed twice and then resuspended in 2% FBS/PBS and 2mM EDTA. Viability dye 7-AAD (1:250; BioLegend, 420403) was added for 10 min at RT. Cells were analyzed with BD FACS Aria Fusion. Mean fluorescence intensity of all viable cells was analyzed using BD FACS DIVA software.

### 20. Bioenergetics

Oxygen consumption rates (OCR) and extracellular acidification rates (ECAR) were measured using Seahorse XFe96 Analyzer (Agilent) and mitochondrial stress test was performed as previously described^97^. Organoids were imaged before the assay, total area determined using the OrgaQuant tool^92^ and then used for normalization.

### 21. Bulk RNA sequencing

Total RNA was isolated from stimulated organoids using RNeasy Micro Kit (Qiagen) and quantified using Qubit High Sensitivity Assay (Invitrogen).

### 22. Sequencing

The process of bulk RNA sequencing was carried out at Novogene using an Illumina Novaseq HiSeq Pair-Ended 150bp, with a sequencing depth of 9G, equivalent to 30 million reads for each sample. Similarly, single-cell RNA sequencing (scRNAseq) was executed, achieving a sequencing depth of 90G for every sample.

### 23. Bulk RNA-seq data analysis

RNA-seq data was preprocessed using the nf-core/rnaseq pipeline (version 3.8.1) ^98^. In brief, reads were aligned using STAR ^99^ to reference genome version GRCh38 with genome annotation version GRCh38 (release 106, Ensembl), and gene expression was quantified using Salmon ^100^.

Differentially expressed genes were identified using DESeq2 ^101^. Genes were included in differential expression analysis if they were detected with more than 1 count-per-million in at least n samples, where n is the number of samples in the smallest group of samples in the comparison. An adjusted p-value (FDR) < 0.1 was applied as threshold to consider genes significantly differentially expressed.

Differential expression of organoids treated with SLE or Control serum: If samples from multiple organoid donors were available, organoid donor was included in the generalized linear model in addition to serum treatment as explanatory variable, thereby accounting for variation from different organoid donors. Contrasts were extracted for serum treatment as the main effect of interest (SLE vs. Control).

Differential expression of human gut epithelial cells of SLE or Control subjects: Gender was included in the generalized linear model in addition to disease status as explanatory variable, thereby accounting for variation from gender. Contrasts were extracted for disease status as the main effect of interest (SLE vs. Control).

Gene set enrichment: Enrichment of terms was calculated using a hypergeometric test (hypeR package ^102^), using gene sets available from the Molecular Signatures Database (MSigDB ^103, 104^). Terms were considered significantly enriched with an adjusted p-value (FDR) < 0.05 threshold. To visualize the similarity of terms based on shared genes, a distance matrix was calculated containing the pairwise distances between terms. As distance metric, binary distance was used, i.e., the proportion of non-shared genes amongst the union of genes from a given pair of terms. The resulting distance matrix was used as input for multidimensional scaling to visualize the similarity of terms in two dimensions.

### 24. IFN score calculation

IFN score was calculated as z-score-based standardized score ^109^, using the transformed count data obtained through variance stabilizing transformation (DESeq2) as input and the following genes as interferon response genes (IRGs): ADAR, BST2, HLA-A, HLA-B, HLA-C, IFI27, IFI35, IFI6, IFIT1, IFIT3, IFIT5, IFITM1, IFITM3, IRF7, IRF9, ISG15, MX1, MX2, OAS1, OAS2, OAS3, OASL, PSMB8, SAMHD1, STAT1, USP18, XAF1.

### 25. Single cell sequencing

Stimulated organoids from eight wells were pooled and dissociated into single cells, bigger fragments removed by a 30 µm strainer and 6000 cells processed using the ChromiumNext GEMSingle Cell 3ʹReagent Kits Dual Index Kit (10x Genomics). The concentration of the prepared library was analyzed, and fragment size confirmed using using High Sensitivity DNA Kit (Agilent).

### 26. Single-cell RNA-seq data analysis

Single-cell RNA-seq data was preprocessed using the nf-core/scrnaseq pipeline (version 2.2.0) ^105^. In brief, read alignment, filtering and counting was performed using cellranger using a reference package created from reference genome version GRCh38 with genome annotation version GRCh38 (release 106, Ensembl). Further analysis steps were performed using Seurat ^106^. Genes that were detected in fewer than 4 cells (from the whole dataset consisting of 34255 cells) were excluded. Cells were included in further analysis if the number of detectable genes was more than 200, the percentage of mitochondrial reads was less than 25%, and the percentage of ribosomal reads was more than 5%. Doublets were excluded using DoubletFinder ^107^.

For further downstream analyses using the Principal Components (e.g., UMAP, nearest-neighbor graph construction for clustering), the first 60 Principal Components were considered. For further downstream analyses with a parameter for number of nearest neighbors (e.g., UMAP, nearest-neighbor graph construction for clustering), a value of 300 and 50 was used for the analysis on all cells and Goblet cells, respectively. For clustering, a cluster resolution of 1 and 0.5 was used for the analysis on all cells and Goblet cells, respectively. For differential expression between two groups of cells, Wilcoxon Rank Sum test was used and genes with an adjusted p-value < 0.1 (Bonferroni correction) were called significant. Trajectory analysis was performed using the Slingshot package^108^ with cluster 0 set to be the cluster of origin (parameter “start.clus”).

### 27. Statistical analyses

All statistical analysis not connected to bulk or single-cell RNA sequencing were performed in GraphPad Prism 9 Software (GraphPad Software, Inc.). Analyses were performed with unpaired t-test or ANOVA with Holm-Šidák’s multiple comparisons test. In the case of organoid size an average of 100 organoids was calculated and the resulting values were compared using chi-square test. To calculate statistical significance for the cytokine measurements undetected cytokines were considered as half of the detection limit. If not stated with exact p values, p values below 0.05 were considered significant and mean with standard deviation is shown

## Discussion

The vital function of the intestinal epithelial cells is to form a barrier that enables tightly regulated passage of a variety of molecules. The barrier integrity highly depends on the presence of mucus bilayer consisting of mucin, mucus-associated proteins and antimicrobial peptides.^110^ Lack of the core mucus protein MUC2 leads to spontaneous development of inflammation in colon and alterations in the mucus layer are associated with a dysfunctional intestinal barrier.^13, 111–114^ Recent reports show an increased intestinal permeability in SLE patients.^15, 23–26^ However, not much is known about the cause and consequence of this observation. Systemic inflammation is marked by hyperactivity of the immune system which is reflected in an altered serum cytokine profile. We theorize that the constant presence of proinflammatory cytokines in the bloodstream and influx to the intestine not only impacts the immune cells in the lamina propria, but it also prompts changes in the epithelial cells. We implemented a model that stimulates organoids with serum to comprehend the primary effects of serum-contained factors on the constantly regenerating epithelial barrier.

With our single cell sequencing analysis, we could observe the presence of nine distinct cell populations. Amongst them were the main cell types of the colon, intestinal stem cells, TA cells as well as cells of the secretory and absorptive lineage. Within the goblet cell population, we could identify not only different goblet cell types with specific function but also a small subfraction of enteroendocrine and Tuft cells. The absorptive lineage was composed of three distinct population whereof one could be assigned to colonocyte progenitors (eCL) and the other two as differentiated colonocytes. Among these two distinct clusters, we discovered a canonical colonocyte population that was highlighted by the expression of markers such as SLC26A3, CA2, and FABP1. Interestingly, the second colonocyte population was marked by a unique expression profile with increased expression of REG4, MMP7, LYZ and the cytokines IL18, IL23, IL33 (Suppl. Table 10). For these genes it has been shown that they can be induced by pro-inflammatory cytokines *in vitro* or *in vivo*. ^12, 80, 82, 115–118^ In addition, we could observe the presence of interferon inducible genes IFI27, ISG15 and ISG20. These findings indicate that the expression profile of this non-canonical colonocyte population was shaped by the cytokine cocktail contained in the serum and identifies a cell population with a specialized function connect to immune defense. Deeper analysis is needed to classify the relevance and function of this cell type *in vivo*. Further studies will be required to fully understand the implications of these findings and their possible relevance in the generation of complex *in vitro* models. Nevertheless, given that most cell types of the descending colon were represented in our model, we could utilize it to study the effects of SLE on the intestinal epithelial barrier.

SLE is a multifaceted autoimmune disease which central player is IFN-α^119–121^. The induction of IFNSG in immune cells has been used as a reliable tool to identify increased serum IFN-α levels as it is the case for the majority of the patients.^46^ Our study shows that serum of SLE patients is able to induce a similar IFNSG in epithelial cells *in vitro*. Thus, intestinal organoids represent a novel tool to assess type-I IFN levels in patient serum. Additionally, we could show that changes induced on the epithelial cells are dependent on type I IFN signaling and can be inhibited by the co-stimulation with anifrolumab an inhibitor of IFNAR1. We further confirmed type I interferon dependency by the absence of DEGs after the stimulation of organoids with serum derived from Granulomatosis with Polyangiitis patients. However, IFN-α in combination with control serum was not able to reproduce the phenotype induced by SLE serum indicating a synergistic effect of type I IFN with other cytokines contained in SLE serum. IL-6, IL-10, IL-18 and IL-23 levels were elevated, and their receptors expressed by the organoids, hence the observed response reflects most likely the integration of all their effects. It was impressive to see that cytokines with a concentration <3 pg/ml were able to induce such a pronounced phenotype demonstrating their potency. In addition, this underlines the artificial environment created using high levels of a cytokine or even a cytokine cocktail for stimulation questioning their relevance in modeling pathological conditions *in vitro*. While they have their relevance in understanding underlying signaling pathways, novel near-physiologic *in vitro* models like the one developed in this study will be of high importance in the future to mimic complex diseases. Our model provides intriguing avenues for exploration into the pathogenesis of SLE.

So far the focus of SLE research has been the dysregulation of the immune system, but studies in mouse intestine (inclusive our own non-published data) and lung cells show that epithelial cells can be affected as well by type I IFN leading to detrimental changes in the barrier.^54, 55^ We could show here that this holds also true for human intestinal cells. Additionally, we observed that mitochondrial pathways and function in the intestinal organoids were affected. SLE serum stimulation caused an increased metabolic activity marked by higher basal respiration and reduced relative spare respiratory capacity highlighting the capacity of serum to induce metabolic changes. A similar mitochondrial dysfunction was recently also reported in CD8+ T cells derived from SLE patients.^46^ In addition, our data suggested that the alteration induced by SLE serum was different from the metabolic shift induced by differentiation. We could see that differentiation led to an increase of oxidative phosphorylation, even in the absence of external energy source promoting this shift *in vivo*^52, 71, 122^. In contrast SLE serum stimulation induced only a slight increase in oxidative phosphorylation but was more dominated by an increase in glycolysis. In UC patients a similar shift towards increased glycolytic activity is accompanied by an impaired goblet cell differentiation. In contrast SLE serum stimulation induced only a slight increase in oxidative phosphorylation but was more dominated by an increase in glycolysis. This shift could mark an overall alteration of mitochondrial function or might reflect the altered cell type composition of the organoids. Since in UC patients a similar shift towards increased glycolytic activity is accompanied by an impaired goblet cell differentiation^71^. Thus, this shift could mark an overall alteration of mitochondrial function or might reflect the altered cell type composition identified upon SLE serum stimulation.

We were intrigued to see that the most common alteration which was persistent in all our experiments was the involvement of the secretory cell lineage. The downregulation of several markers connected to secretion, goblet and enteroendocrine cells identified with bulk RNA sequencing aligned with our results from microscopy and single-cell RNA sequencing which revealed a diminished goblet and enteroendocrine cellpopulation and a decrease or delay in the differentiation towards secretory cells. Additionally, our results indicated a functional change of differentiated cells resulting in an altered mucus layer. Those changes were not restricted to the goblet cell population but extended to the absorptive cell lineage. We classified numerous misregulated genes, many of which were integral to mucus composition (AGR2, SPINK4, MUC12, FCGBP) and antimicrobial defense (PLA2G2A, RETNLB, REG4, TFF3, and WFDC2). Our data aligned with the altered expression pattern of several mucus related genes and the reported metaplastic, colonic expression of LYZ associated with UC pathogenesis^12, 13, 65, 123–125^. We observed downregulation of MUC12, a membrane-bound mucin that has a protective role for colonocytes.^126^ FCGBP, important for mucus integrity^12, 123, 124, 127^, and MMP7 which is associated with LPS-induced barrier permeability and connected to SLE pathogenesis^81, 128^ itself were upregulated. The additionally identified downregulation of the sodium channel SCNN1A and the aquaporin AQP5 suggests an alteration in ion composition and hydration of the mucus^57, 129^. With scRNA-seq we were able to identify downregulation of tight junction proteins CLDN3, CLDN7 and TJP1 that indicate a change in intestinal epithelial permeability. Furthermore, we could show that non-canonical colonocytes and a subcluster of goblet cells were specifically responsive to IFN contained in the serum by the expression of IFI27 indicating a cell type specific response.

These findings show that SLE serum stimulation led to an alteration in both the secretory and absorptive lineage that resulted in a potential structural weaking of the mucus barrier. Not only the differentiation to goblet cells and therewith a reduction in their number was affected but also several secreted factors necessary for a functional mucus layer. The alteration in expression of AMPs suggested changes in intestinal antimicrobial capacity of the mucus which is pivotal to maintain the sterility of the inner mucus layer and therewith reducing the microbial-epithelial interaction *in vivo*.^130^ It is important to mention that in UC it is suggested that mucus alterations are causative for disease pathogenesis and precede the active disease, indicating the importance of the mucus layer for barrier function and intestinal homeostasis.^13^ Also other diseases like cystic fibrosis highlight the importance of a physiological mucus composition in order to maintain organ function and underline that even minor changes to the mucus layer can have potentially detrimental effects on barrier funtion.^131^ Additional studies are required to confirm changes in the physical properties of the mucus layer in our model system and to understand how these changes might impact the epithelial barrier *in vivo*. It would be of specific interest to unravel if changes in the mucus layer could trigger changes in microbiome composition reported for SLE^132–134^.

The relevance of the changes unraveled with our *in vitro* model was highlighted by the findings revealed in a small SLE patient population. We observed that in two of three analyzed descending colon samples we could detect a clear effect on the epithelial cells manifested by misregulation of ion transporters SLC26A3, SCNN1A and SCNN1B expression along with BEST2, a goblet cell marker, and the reduced expression of several other goblet cell related markers. It is of high interest to understand if initiated by these changes mucus layer composition and function are altered. A bigger and well-stratified patient population with matched controls is necessary to confirm these initial findings that point towards an involvement of the intestinal epithelial cells in SLE pathogenesis.

To conclude, our research underscores the immense potential of organoids in translational research, offering a fresh perspective to study diseases that have stumped medical science for decades. The adaptability of organoids permits the incorporation of different factors, such as disease-specific cytokines or even other cell types, further emphasizing their potential in advancing medical research.

## Figure Legends

**Suppl. Fig1:** A: Fluorescence microscopy images of intestinal colon organoids stimulated with control (top panel) or SLE (bottom panel) serum revealing no differences in apoptosis (cleaved Caspase 3 staining) after 72h serum stimulation. Scale bar 100 µm. (B) Heatmap shows cell type composition variation in both organoid lines by various cell type markers for proliferation and differentiation due to individual proliferation and differentiation dynamics, n=10 organoid wells per organoid line.

**Suppl. Fig 2:** A. Heatmap visualizing the expression of selected IFN-inducible genes, showing an upregulation across all SLE serum-stimulated organoids from line I. n= 10 control serum (samples a to j) and n=8 SLE serum (samples 1 to 8). B. Volcano plot showing the most significantly up- or downregulated genes in IFN-α-serum compared to additional anifrolumab treatment. Heatmap demonstrating gene expression profiles of organoids treated with control serum plus IFN-α in the presence or absence of anifrolumab. Genes relevant in IFNα/β signaling are displayed, and 27 specific genes were identified as unique to the epithelial cell response to a low dose of IFN-α, which were completely downregulated when blocking IFNAR1 with anifrolumab. n=3 IFNα+control serum, and n=3 IFNα+control serum+anifrolumab. C. Graph depicting normalized counts for IFNα-1 and IFNα-2 in serum stimulated organoids of line I and II. Absence of main interferon transcripts validates that the observed IFN signature was not due to endogenous IFN expression. n= 10 control serum (samples a to j) and n=8 SLE serum (samples 1 to 8). D. Western blot illustrating the levels of STAT1 and IFIT3 in organoids from SLE and control conditions after 48h of stimulation at two different serum concentrations (5% and 30%). In the IFNα-stimulated organoids, STAT1 and IFIT3 levels were assessed after 1h and 48h of stimulation at a concentration of 100ng/ml. Each sample represents two pooled wells of organoids stimulated with serum from one donor.

**Suppl. Fig. 3:** A. Heatmap demonstrating upregulation of mitochondrial genes in organoids stimulated with SLE serum, based on expression levels across various serum donors on organoid line II. n= 10 control serum (samples a to j) and n=8 SLE serum (samples 1 to 8). B. Graphical representation of Mitored measured by flow cytometry data displaying no significant changes in mitochondrial mass between organoids stimulated with SLE serum and control serum. Each dot represents one well of organoids dissociated into single cells and analyzed for mean fluorescence intensity (MFI); n=4 for each condition. C. Seahorse assay revealing increased ATP production and no changes in maximal respiration in SLE serum stimulated organoids. Each dot corresponds to one well of organoids stimulated with n=10 control serum samples or n=8 SLE serum samples. Each serum stimulation was analyzed with two technical replicates. D. Seahorse assay comparing organoids with different cell differentiation states, revealing decreased spare respiratory capacity in differentiated organoids. Each dot corresponds to one well of organoids; n=12 wells per condition. E. Energy map plotted from Seahorse assay comparing organoids with different cell differentiation states. PRO: Organoids with cells in proliferative state; DIF: Organoids containing also differentiated cells. Statistical tests performed with unpaired t-test.

**Suppl. Fig. 4:** A. Graphical representation and corresponding fluorescence microscope images of the proliferative capability of organoids following SLE serum stimulation, as measured by the ratio of Ki67-positive cells to total cell nuclei (Scale bar: 50 µm). Each dot in the graph represents one organoid, n=98 organoids stimulated with control or n=78 SLE serum No significant differences in proliferative capacity between organoids stimulated with SLE serum and control serum was observed as analyzed by unpaired t-test. B. Graphical representation and corresponding fluorescence microscope images showing S phase progression measured by EdU incorporation per cell relative to the total cell count (Scale bar: 100 µm). Each dot in the graph represents one organoid. Comparable S phase dynamics are observed between SLE serum-stimulated organoids and control conditions as analyzed by unpaired t-test. C. Heatmap illustrating downregulated genes associated with the term secretory vesicle and mucus in 72h SLE serum stimulated organoids in organoid line I; n= 10 control serum (samples a to j) and n=8 SLE serum (samples 1 to 8).

**Suppl. Fig5:** A. Western blot depicting phosphorylation of STAT1 to confirm the inhibitory effect of anifrolumab at varying concentrations in response to IFNα2 stimulation in organoids. B. Graphical representation of the differential impact on relative spare respiratory capacity in organoids following exposure to serum from control (n=10), SLE (n=10), unstimulated control (NC) (n=3), and granulomatosis with polyangiitis (GPA) patients (n=3); two technical replicates except for NC are depicted. The results demonstrate that the mitochondrial dysfunction observed in organoids treated with SLE patient serum is not observed in those exposed to GPA patient serum. Each dot represents one well of organoids. Statistical test performed with one way ANOVA with Holm-Šidák’s multiple comparisons test. C. Venn diagram showing the overlap of DEGs resulting from stimulation with SLE compared to control serum and IFNα+control serum compared to control serum. The small overlap is mostly constituted by IFN-related genes, highlighting the unique response of organoids to SLE serum, which is not purely induced by IFN but other SLE-specific serum components. Statistical test performed with one way ANOVA with Holm-Šidák’s multiple comparisons test.

**Suppl. Fig.6:** A. UMAP demonstrates the distribution of all examined cells derived from single-cell RNA-Seq data per sample. These cells come from control (n=3) and SLE (n=3) serum-stimulated organoids and are color-coded to represent 9 distinctive clusters. B. Heatmap representing the top 20 DEG per identified cluster. C: Diagram representing the four different predicted cellular developmental trajectory using Slingshot. D: Uniform Manifold Approximation and Projection (UMAP) demonstrates the distribution of all examined cells derived from single-cell RNA-Seq data per sample within the goblet cell subclusters. E. UMAP representation of CHGA and POU2F3 expression level in goblet cells subclusters. F. Diagram representing the predicted cellular developmental trajectory within the goblet cell subcluster using Slingshot.

**Suppl. Fig. 7. A:** Dot plot depicting average expression of ribosomal genes post SLE serum stimulation for each cell cluster. The size and color of each dot represent the cell percentage and average gene expression, respectively. B: Dot plot depicting average expression of oxidative phosphorylation genes post SLE serum stimulation for each cell cluster. The size and color of each dot represent the cell percentage and average gene expression, respectively. C: Bar graph showing the distributions of goblet cell subclusters within the total analyzed cells for SLE (n=3) and control (n=3) serum stimulated organoids. Unpaired t-test was performed.

## Acknowledgments

We deeply appreciate the substantial contributions to this study from individuals and institutions alike. First, our sincere thanks to Hugo deJonge for his crucial donation of the Rspondin cell line. We also express our gratitude to Alina Winkelotte and Almut Schulze for their invaluable help with the Seahorse analysis. Stephan Schöler’s expertly crafted data analysis scripts significantly expedited our research and clarified our findings. The quality of our work was greatly enhanced by the critical input and guidance from Christoph Becker and Daigen Xu. Our findings were deeply enriched by the insightful contributions from Alexander Drainas and Mojca Frank-Bertoncelj. Rafael Sênos Demarco provided invaluable advice on mitochondria assays. Our thanks also go to Novogene, with their assistance with RNA sequencing, the Heidelberg Nikon Imaging Center at the University of Heidelberg, part of BioQuant, for providing necessary resources and facilities. Special recognition to the patients, Novabiosis, DTI foundation, and the I3PT Biobank for their collaboration. We particularly thank Joaquim Albiol, Fernando Mosteiro, and his dedicated team for their indispensable help. All contributions, regardless of size, profoundly impacted the success of this study, and we apologize for any unintentional omissions. The research was financially supported by Merck KGaA, Darmstadt, Germany, during the employment of MRD, IH, SE, MS, and BS at the BioMed X Institute.

## 30. Author contributions

**IH:** Conceptualization; data curation; formal analysis; validation; investigation; visualization; writing – original draft; writing – review and editing. **SE:** formal analysis, visualization, review, and editing. **MS:** Investigation. **BS:** Investigation. **SB**: Investigation. **WM:** Resources. **SK:** Resources. **HML:** Resources. **JH:** Resources. **BW:** Resources**. MRD:** Conceptualization; formal analysis; supervision; funding acquisition; investigation; visualization; writing – original draft; project administration; writing – review and editing.

## 31. Data availability

RNA sequencing data that support the findings of this study are in process to be uploaded at European Genome-Phenome Archive (EGA). Analyzed transcriptomic data are as well provided as Supplementary Tables. All other data supporting the findings of this study are available from the corresponding author on request.

