## Supplemental figures for "Systemic Lupus Erythematosus Serum Stimulation of Human Intestinal Organoids Induces Changes in Goblet Cell Differentiation and Mitochondrial Fitness"

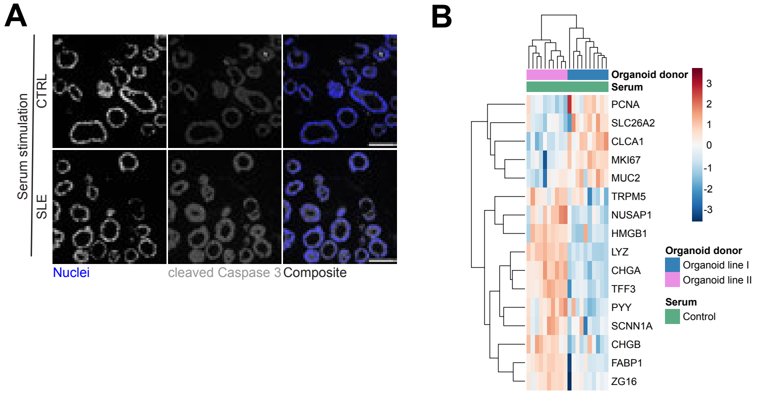


Suppl. Figure 1

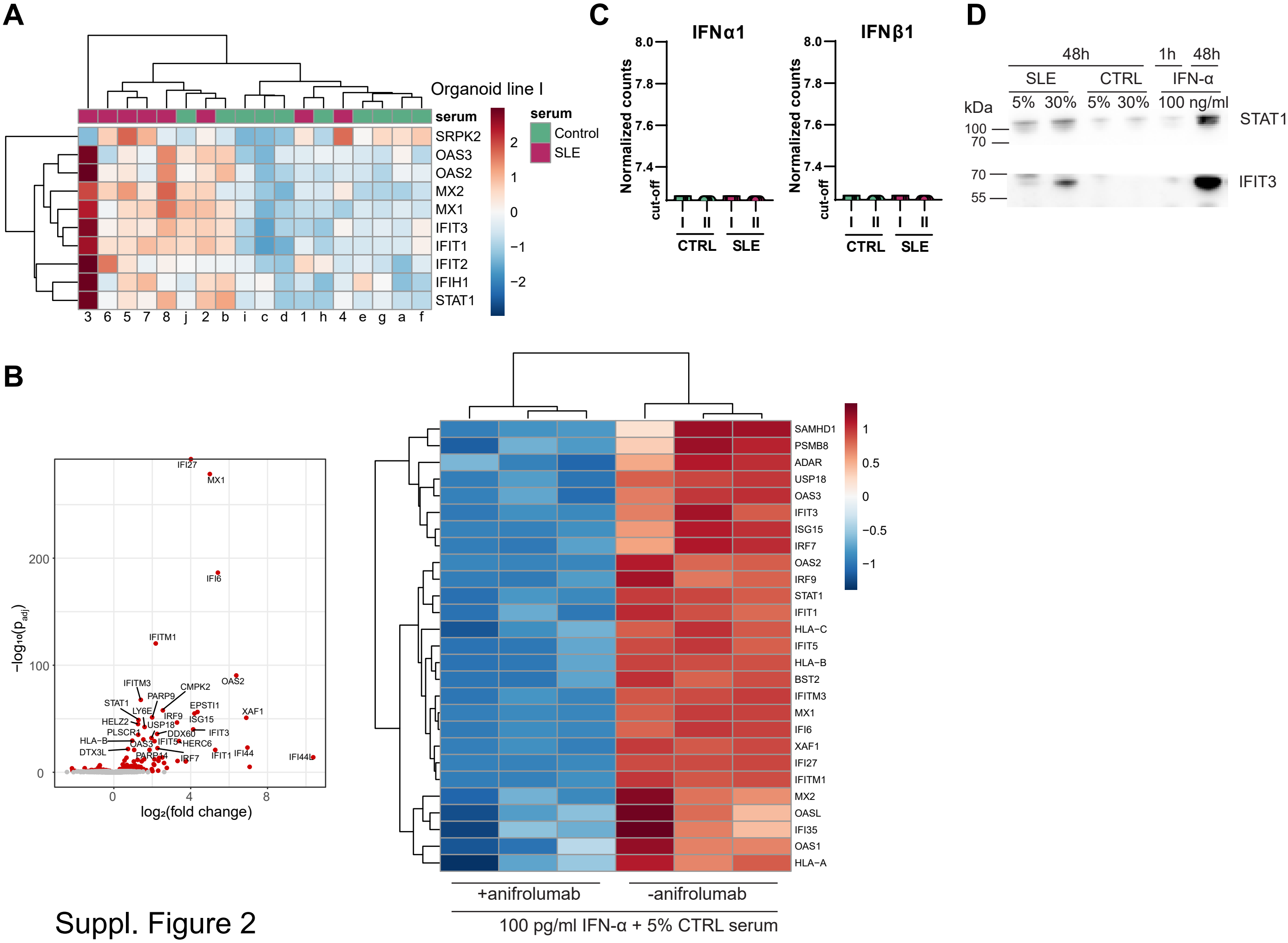


Suppl. Figure 2

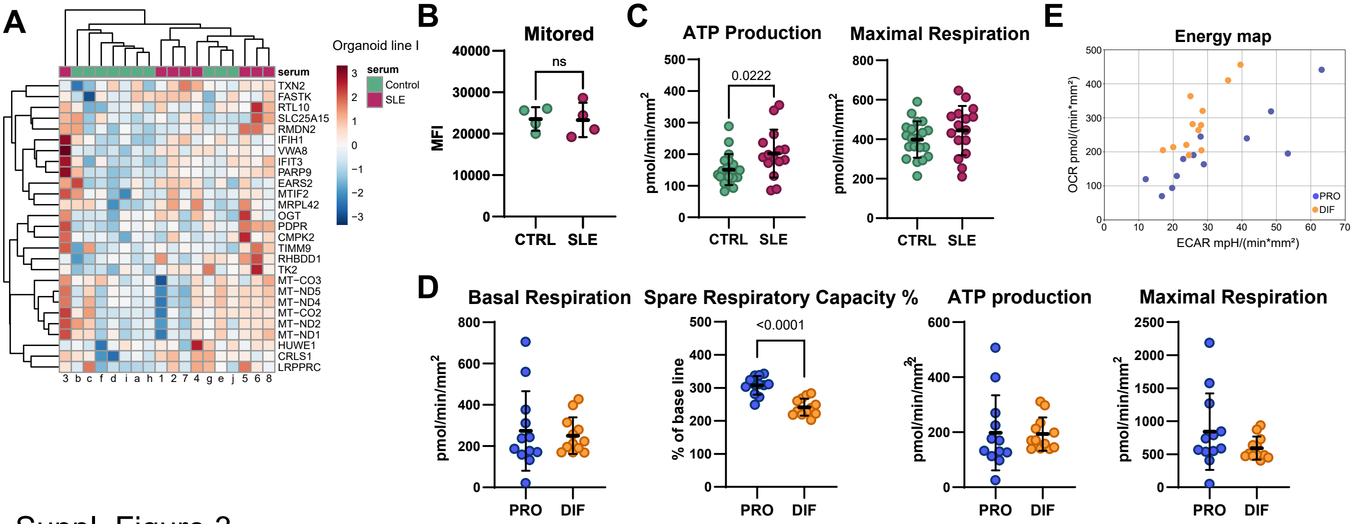


Suppl. Figure 3

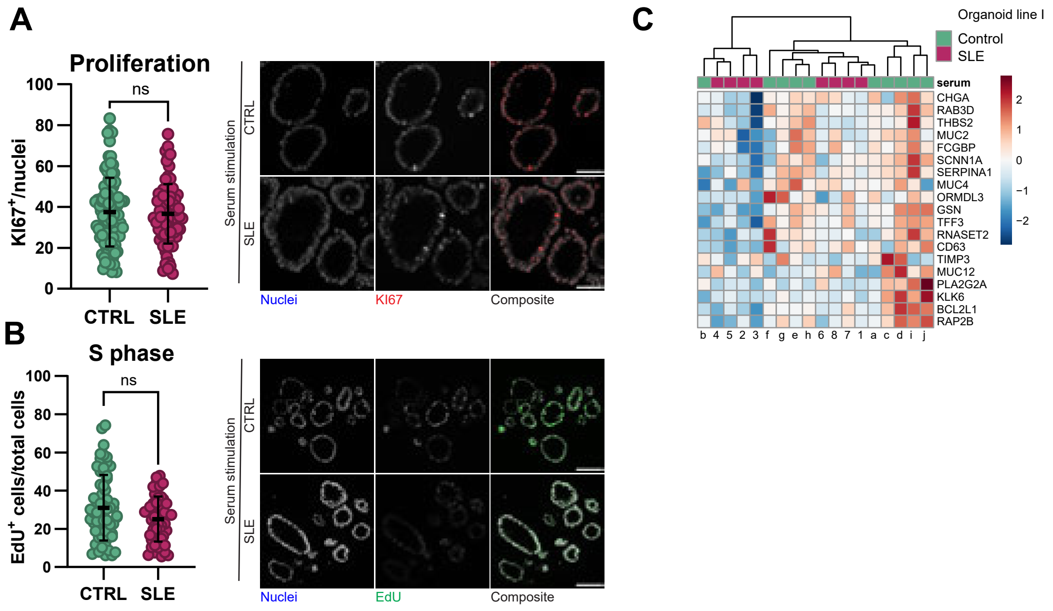


Suppl. Figure 4

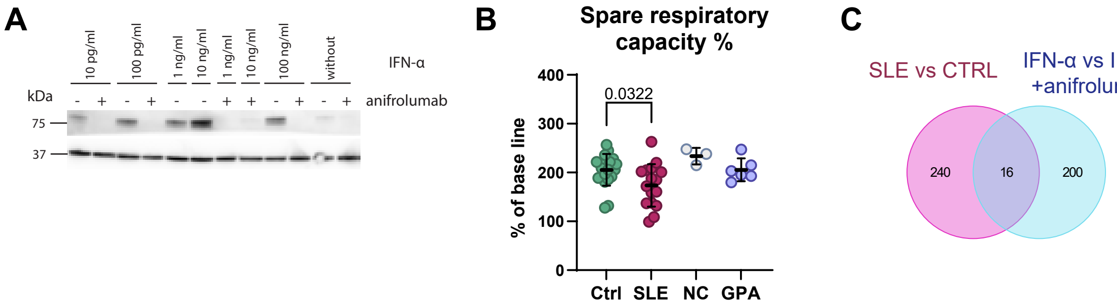


| Overlap | |
|---|---|
| CMPK2 | OAS2 |
| DHX58 | OAS3 |
| EPSTI1 | PARP14 |
| IFIH1 | PARP9 |
| IFIT1 | PHF11 |
| IFIT3 | SAMD9 |
| MX1 | SAMD9L |
| MX2 | SP110 |

Suppl. Figure 5

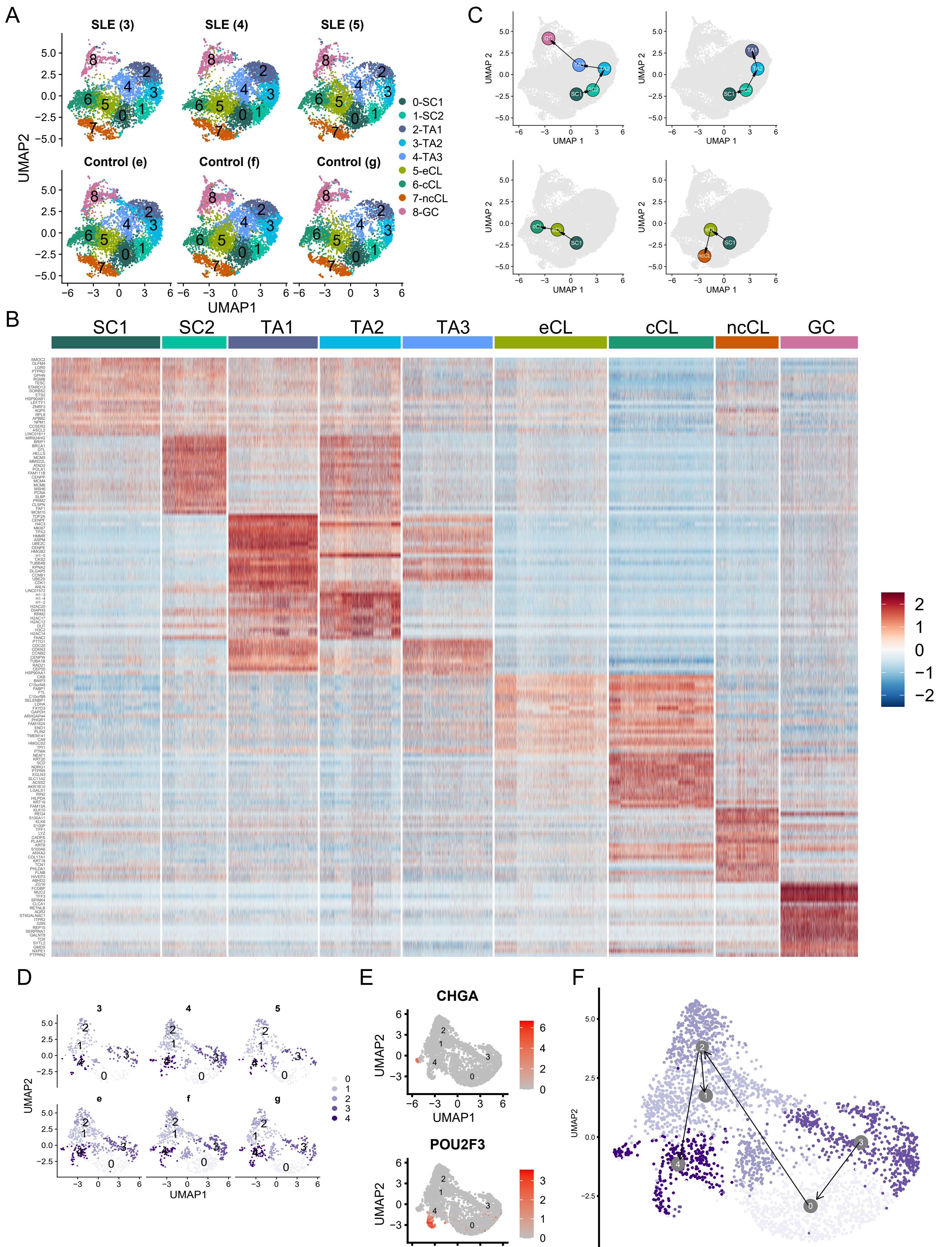


Supplementary Fig. 6

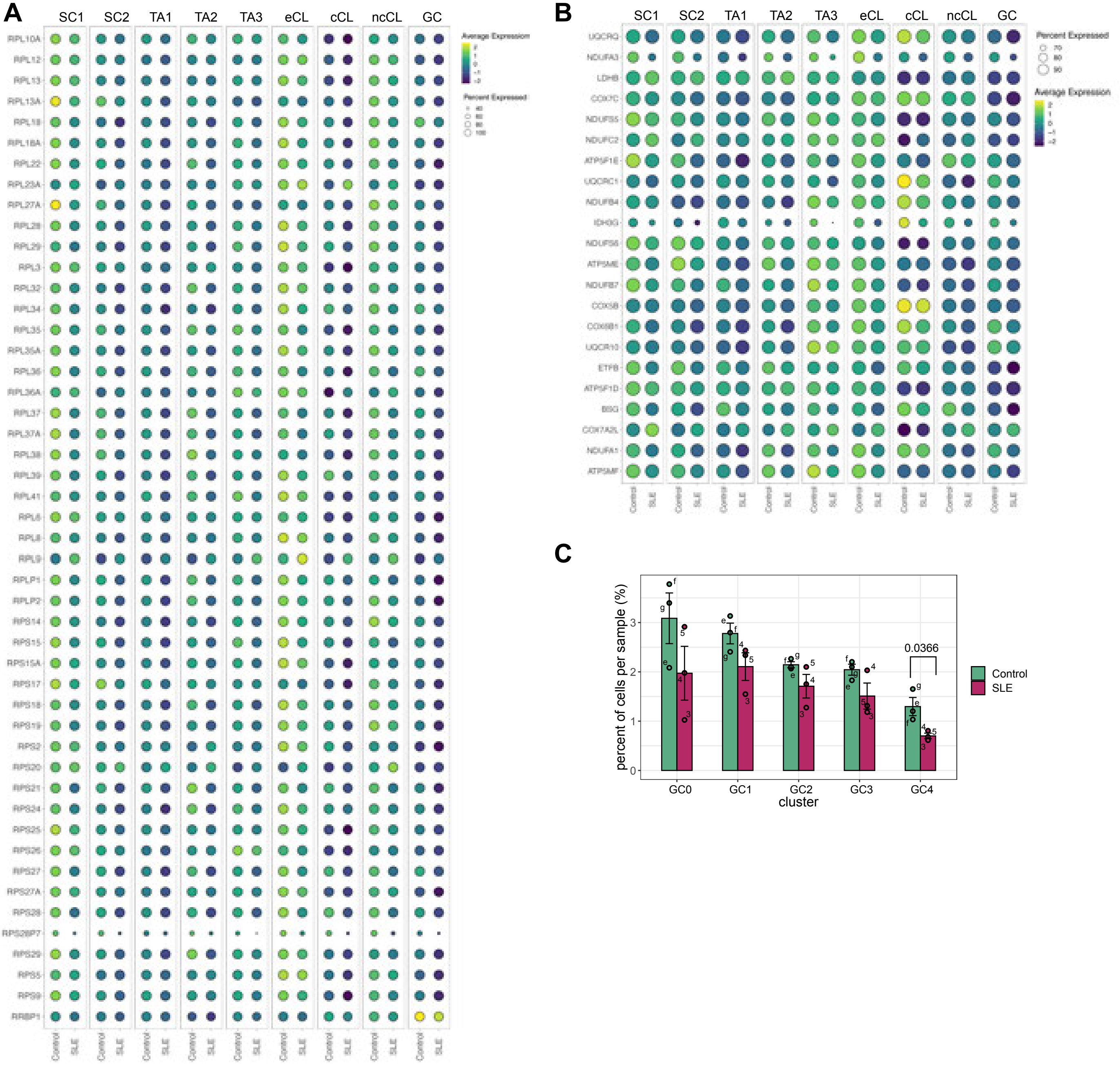
A
SC1
SC2
TA1
TA2
TA3
eCL
cCL
ncCL
GC
Average Expression
Percent Expressed
RPL10A
RPL12
RPL13
RPL13A
RPL18
RPL18A
RPL22
RPL23A
RPL27A
RPL28
RPL29
RPL3
RPL32
RPL34
RPL35
RPL35A
RPL36
RPL36A
RPL37
RPL37A
RPL38
RPL39
RPL41
RPL6
RPL8
RPL9
RPLP1
RPLP2
RPS14
RPS15
RPS15A
RPS17
RPS18
RPS19
RPS2
RPS20
RPS21
RPS24
RPS25
RPS26
RPS27
RPS27A
RPS28
RPS28P7
RPS29
RPS5
RPS9
RRBP1
Control
SLE
B
UQCRQ
NDUFA3
LDHB
COX7C
NDUFS5
NDUFC2
ATP5F1E
UQCRC1
NDUFB4
IDH3G
NDUFS6
ATP5ME
NDUFB7
COX5B
COX6B1
UQCR10
ETFB
ATP5F1D
BSG
COX7A2L
NDUFA1
ATP5MF
C
percent of cells per sample (%)
0.0366
Control
SLE
GC0
GC1
GC2
GC3
GC4
cluster

Supp. Fig. 7
